# Expanding and improving analyses of nucleotide recoding RNA-seq experiments with the EZbakR suite

**DOI:** 10.1101/2024.10.14.617411

**Authors:** Isaac W. Vock, Justin W. Mabin, Martin Machyna, Alexandra Zhang, J. Robert Hogg, Matthew D. Simon

## Abstract

Nucleotide recoding RNA sequencing methods (NR-seq; TimeLapse-seq, SLAM-seq, TUC-seq, etc.) are powerful approaches for assaying transcript population dynamics. In addition, these methods have been extended to probe a host of regulated steps in the RNA life cycle. Current bioinformatic tools significantly constrain analyses of NR-seq data. To address this limitation, we developed EZbakR, an R package to facilitate a more comprehensive set of NR-seq analyses, and fastq2EZbakR, a Snakemake pipeline for flexible preprocessing of NR-seq datasets, collectively referred to as the EZbakR suite. Together, these tools generalize many aspects of the NR-seq analysis workflow. The fastq2EZbakR pipeline can assign reads to a diverse set of genomic features (e.g., genes, exons, splice junctions, etc.), and EZbakR can perform analyses on any combination of these features. EZbakR extends standard NR-seq mutational modeling to support multi-label analyses (e.g., s^4^U and s^6^G dual labeling), and implements an improved hierarchical model to better account for transcript-to-transcript variance in metabolic label incorporation. EZbakR also generalizes dynamical systems modeling of NR-seq data to support analyses of premature mRNA processing and flow between subcellular compartments. Finally, EZbakR implements flexible and well-powered comparative analyses of all estimated parameters via design matrix-specified generalized linear modeling. The EZbakR suite will thus allow researchers to make full, effective use of NR-seq data.

## Introduction

Developing a mechanistic understanding of gene expression regulation requires methods to probe the kinetics of RNA synthesis, processing, transport, and degradation. Standard RNA-seq provides limited information about the kinetics of the processes that determine the abundance of an RNA. Nucleotide recoding RNA-seq (NR-seq; TimeLapse-seq, SLAM-seq, TUC-seq, etc.) overcomes these limitations (Herzog, et al., 2017; Riml, et al., 2017; Schofield, et al., 2018). NR-seq combines metabolic labeling with chemistries that recode the hydrogen bonding pattern of the metabolic label so as to facilitate detection of labeled RNA via chemically induced mutations in sequencing reads, without the need for biochemical enrichment of the labeled RNA. By providing information about both overall RNA abundance and the dynamics of nascent and pre-existing RNA, NR-seq resolves the kinetic ambiguities of standard RNA-seq (Duffy, et al., 2019; Erhard, et al., 2022).

The field has seen an explosion of methodologies using NR-seq to study different aspects of the RNA life cycle. For example, transient transcriptome sequencing (Schwalb, et al., 2016) has been extended to improve studies of nascent and short-lived RNAs (Reichholf, et al., 2019; Schofield, et al., 2018). Similarly, nucleotide recoding has been combined with Start-seq to study the kinetics of promoter-proximal pausing (Nechaev, et al., 2010; Zimmer, et al., 2021), Ribo-seq to study the kinetics of translation initiation and elongation (Ingolia, et al., 2009; Schott, et al., 2021), PacBio long read sequencing to improve studies of transcript isoform dynamics (Rahmanian, et al., 2020; Rhoads and Au, 2015), subcellular fractionation to better understand RNA flow between subcellular compartments (Ietswaart, et al., 2024; Müller, et al., 2024; Steinbrecht, et al., 2024), and tethering experiments to probe small molecule binding sites in the transcriptome (Moon, et al., 2024). Nucleotide recoding has also been combined with a number of scRNA-seq protocols to facilitate probing synthesis and degradation kinetics in individual cells (Cao, et al., 2020; Erhard, et al., 2019; Hendriks, et al., 2019; Lin, et al., 2023; Maizels, et al., 2024; Qiu, et al., 2020) and promises to significantly improve analyses of RNA velocity by providing an alternative to using the potentially biased or non-existent intronic coverage at key regulatory genes (Maizels, et al., 2024; Qiu, et al., 2022; Weiler, et al., 2024). Finally, while nucleotide recoding chemistries were originally developed for 4-thiouridine (s^4^U), some have been shown to successfully recode 6-thioguanosine (s^6^G) as well (Gasser, et al., 2020; Kiefer, et al., 2018). This has opened the door for multi-label experimental designs (Courvan, et al., 2022). This family of NR-seq extensions has already provided a host of novel biological insights, and its continued growth creates a need for better and more accessible analysis tools for NR-seq data to expand our understanding of RNA biology.

We aimed to develop a suite of tools that can:

1. Analyze reads assigned to a flexible array of genomic features (e.g., exons, introns, exonic bins, splice junctions, transcript equivalence classes, etc.). This facilitates integration with differential expression analyses capable of working with the same feature sets and allows users to make full use of all of their aligned NR-seq reads.
2. Make the processed mutational data readily available to users to support assessment of model fits and future model improvements.
3. Implement a rigorous statistical model of the mutational data in NR-seq reads to estimate the abundances of labeled and unlabeled RNA. These models should also be compatible with multi-label data.
4. Support flexible kinetic modeling that is compatible with standard analyses of mature total RNA as well as analyses of premature mRNA dynamics and flow between subcellular compartments to better support the full range of NR-seq methods.
5. Perform well-powered and flexible comparative analyses of any estimated kinetic parameter to enable in-depth investigations of experimental perturbations.

To date, no existing tools have all these capabilities. pulseR, originally designed for enrichment-based methods, has been repurposed for NR-seq data, but may not be suitable for enrichment-free data, and was not designed to process NR-seq data (Uvarovskii and Dieterich, 2017). SLAMDUNK, one of the first pipelines developed to process raw NR-seq data, is widely used but was specifically optimized for single-end, 3’-end sequencing data (Herzog, et al., 2017; Neumann, et al., 2019). In addition, it uses a mutation count cutoff to distinguish labeled and unlabeled reads, which can yield inaccurate kinetic parameter estimates that are dependent on the metabolic label incorporation rate in a particular sample. A gold-standard analysis pipeline must overcome the limitations of this labeled vs. unlabeled read classification approach.

Our lab developed a two-component mixture model to estimate the fraction of sequencing reads derived from labeled RNA (Schofield, et al., 2018). This strategy was subsequently implemented in the user-friendly and performant software tool GRAND-SLAM (Jürges, et al., 2018). GRAND-SLAM performs analyses at the gene-level, allowing users to either retain or throw out reads mapping to intronic regions. Separately analyzing premature and mature mRNA dynamics with GRAND-SLAM is challenging and requires custom workarounds (Ietswaart, et al., 2024). In addition, GRAND-SLAM provides mixture model fits as output, but provides limited access to the intermediate mutational data passed to the mixture model. Providing such data in a convenient format would help users explore their mutational data and facilitate development of improved modeling strategies (Ietswaart, et al., 2024; McShane, et al., 2024; Müller, et al., 2024; Steinbrecht, et al., 2024). Finally, neither GRAND-SLAM nor the recently developed helper R package grandR (Rummel, et al., 2023) support well-powered comparative analyses akin to those implemented in differential expression analysis software (Love, et al., 2014; Robinson, et al., 2010).

We previously addressed some of these limitations with the development of a pipeline (bam2bakR) and R package (bakR) (Vock and Simon, 2023). bam2bakR provides as output a compressed representation of the processed mutational data, which can be analyzed via mixture models implemented in bakR. In addition, bakR borrows strategies from differential expression analysis tools to perform well-powered comparative analyses of kinetic parameters and better support downstream biological investigations (Love, et al., 2014; Robinson, et al., 2010). While bam2bakR can separately assign reads to exonic and intronic regions, bakR provides limited support for making use of that information to study pre-mRNA dynamics, as it can only perform analyses on one feature set at a time (e.g., exonic regions of genes). bakR’s rigid design and lack of modularity also limits the type of NR-seq datasets it can analyze. Finally, no existing tool supports analyses of multi-label experiments or flexible dynamical systems modeling of NR-seq data. The field needs further bioinformatic innovation to make more effective use of NR-seq data.

Here we present the EZbakR suite, a Snakemake pipeline (fastq2EZbakR) and R package (EZbakR) designed to support flexible analyses of NR-seq experiments (Figure 1). fastq2EZbakR is able to assign reads to a wide array of genomic features, and EZbakR supports analyses of any combination of these features. EZbakR generalizes the mutational mixture model implemented in GRAND-SLAM and bakR to support multi-label designs. It also optionally implements an improved hierarchical model that allows individual features to have unique incorporation rate estimates, addressing a recently discovered source of variance in NR-seq data (McShane, et al., 2024). In addition, EZbakR generalizes fitting models of linear dynamical systems to NR-seq data so as to support analyses of pre-mRNA dynamics and of subcellular fractionation-based extensions of standard NR-seq (Ietswaart, et al., 2024; Müller, et al., 2024; Steinbrecht, et al., 2024). Finally, EZbakR implements design matrix specified comparative analyses of kinetic parameters to radically increase its flexibility relative to its predecessor, bakR. We use a panel of simulated data to validate all of these novel functionalities and showcase how the EZbakR suite promises to facilitate unprecedented investigations of existing and future NR-seq datasets.

**Figure 1:**
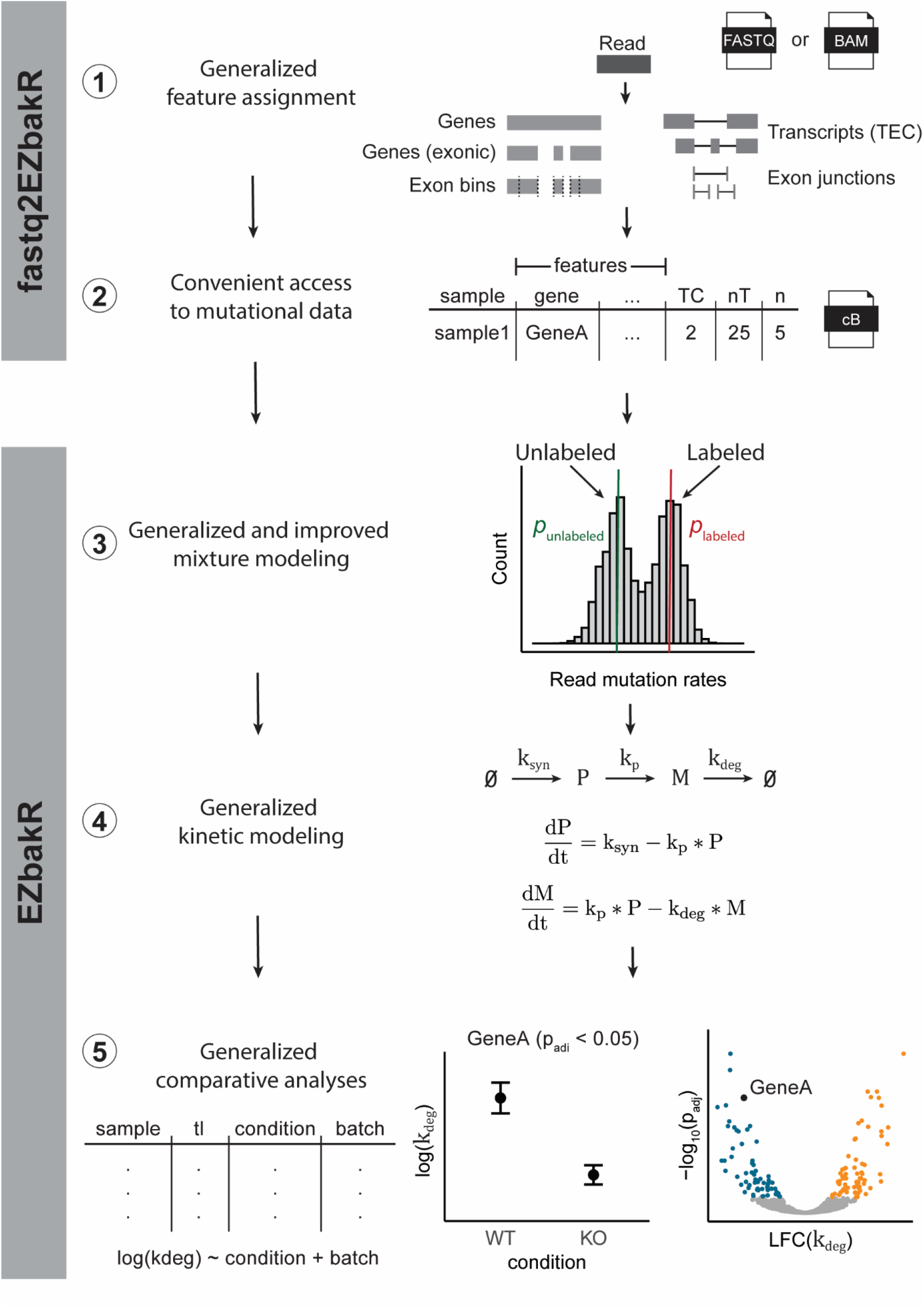
The EZbakR suite generalizes and improves upon all steps of the NR-seq analysis pipeline. The EZbakR suite: 1) Implements a flexible feature assignment strategy, 2) provides processed mutational data in a convenient, compressed format, 3) analyzes mutational data in a way that supports multi-label design and allows for feature-to-feature mutation rate variance, 4) fits any identifiable, linear dynamical systems model to NR-seq data, and 5) performs well-powered, design matrix-specified comparative analyses of all estimated kinetic parameters.

## Results

### Generalized feature assignment

In theory, reads in an NR-seq experiment can be assigned to any annotated genomic feature (e.g., genes), and the fraction of reads derived from labeled RNA can be estimated for each feature. In practice, existing pipelines provide limited flexibility in terms of the types of features to which they assign reads. To address this limitation, fastq2EZbakR supports much more flexible feature assignments than any existing NR-seq analysis pipeline (Figure 2). In particular, reads can be assigned to genes (exons and introns), exclusively exonic regions of genes, exonic bins (Anders, et al., 2012), transcript equivalence classes (TECs; i.e., the set of transcript isoforms with which a read is compatible) (Cmero, et al., 2019), and exon-exon junctions (Figure 2A). fastq2EZbakR also implements the read pre-processing, alignment, and mutation counting necessary for NR-seq data (Figure 2B). The EZbakR input data object is highly flexible (see EZbakR documentation and Figure 1, point 2). EZbakR is able to accept information about read assignment to any arbitrary combination of features, and can perform analyses on any subset of these features (or the full set of features). This both allows analyses performed by EZbakR to be easily integrated with that of expression analysis tools like DEseq2, edgeR, DEXSeq, etc., and opens the door for integration across different feature-level analyses (e.g., pre-mRNA and mature mRNA) (Anders, et al., 2012; Love, et al., 2014; Robinson, et al., 2010). The EZbakR suite thus generalizes feature assignment and analysis of NR-seq reads.

**Figure 2:**
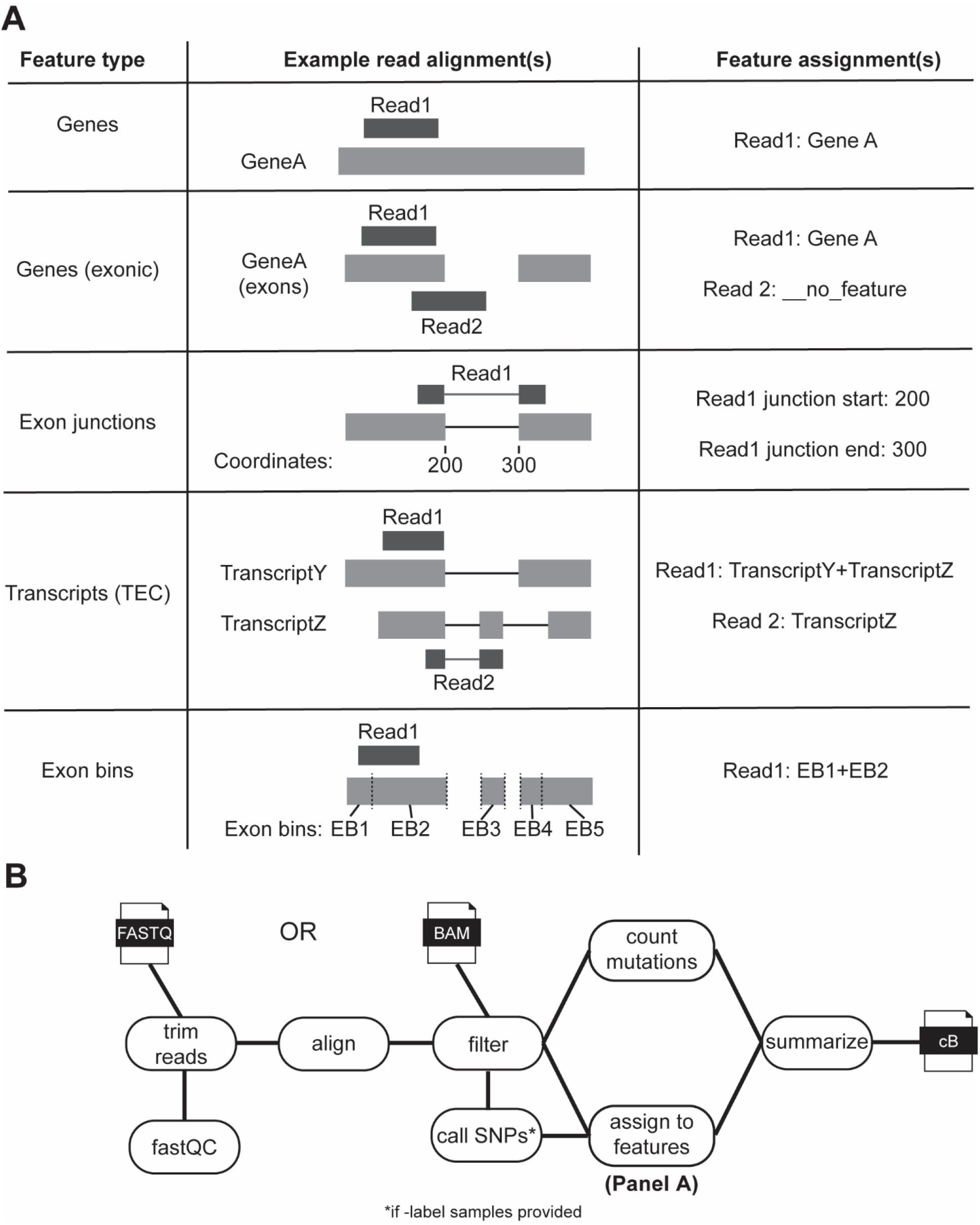
fastq2EZbakR generalizes feature assignment to support finer dissection of NR-seq data. **A)** Schematic of the 5 different feature assignment strategies implemented in fastq2EZbakR. If a read does not overlap with a given feature, it will be assigned a value of no_feature for that assignment. If a read overlaps multiple features, all features will be reported, with names separated by +-signs. TEC = transcript equivalence class (set of transcript isoforms with which a read is compatible). Exon bins were introduced in DEXSeq (Anders et al., 2014). **B)** Schematic of the full fastq2EZbakR pipeline.

### Generalized and improved mixture modeling

After reads have been assigned to features and mutations in each read counted, the next step of any NR-seq analysis is to figure out how many sequencing reads are derived from labeled RNA for each feature (i.e., the fraction labeled, denoted *θ* in figures). While it is common to define a mutation content cutoff such that all reads with that many mutations or more are attributed to labeled RNA, this strategy can yield biased estimates (Figure S1). Mixture modeling is a far more accurate, rigorous, and robust strategy, and has been implemented in several existing tools (Jürges, et al., 2018; Schofield, et al., 2018; Vock and Simon, 2023). In this strategy, assumptions are made regarding distributions that accurately describe the probabilities of seeing N mutations in reads from labeled and unlabeled RNAs with M mutable reference nucleotides. Typically, this means assuming that a read’s mutational content is binomially distributed, conditional on its status as being from labeled or unlabeled RNA. GRAND-SLAM and bakR both use this strategy to analyze single-label NR-seq data. In EZbakR, we have generalized this model to include analyses of multi-label NR-seq datasets (Figure 3A). To showcase this generalized model, we considered one such method, TILAC, where s^4^U labeled cells are mixed with s^6^G labeled cells (Figure 3B) (Courvan, et al., 2022). TILAC allows for internally normalized comparative analyses, analogous to the proteomic method SILAC (Mann, 2006). In TILAC, there are three populations of reads: those with high T-to-C mutation rates, those with high G-to-A mutation rates, and those with background level T-to-C and G-to-A mutation rates (Figure 3C). Reads with both high T-to-C and G-to-A mutation rates cannot exist in this case, as no cell was exposed to both labels. EZbakR is flexible and able to account for the absence of this fourth population (Figure 3C mutational populations table). Analyses of simulated TILAC data confirmed that EZbakR’s multi-label mixture model fit is accurate (Figure 3D). The EZbakR suite thus generalizes mixture modeling of NR-seq data.

**Figure 3:**
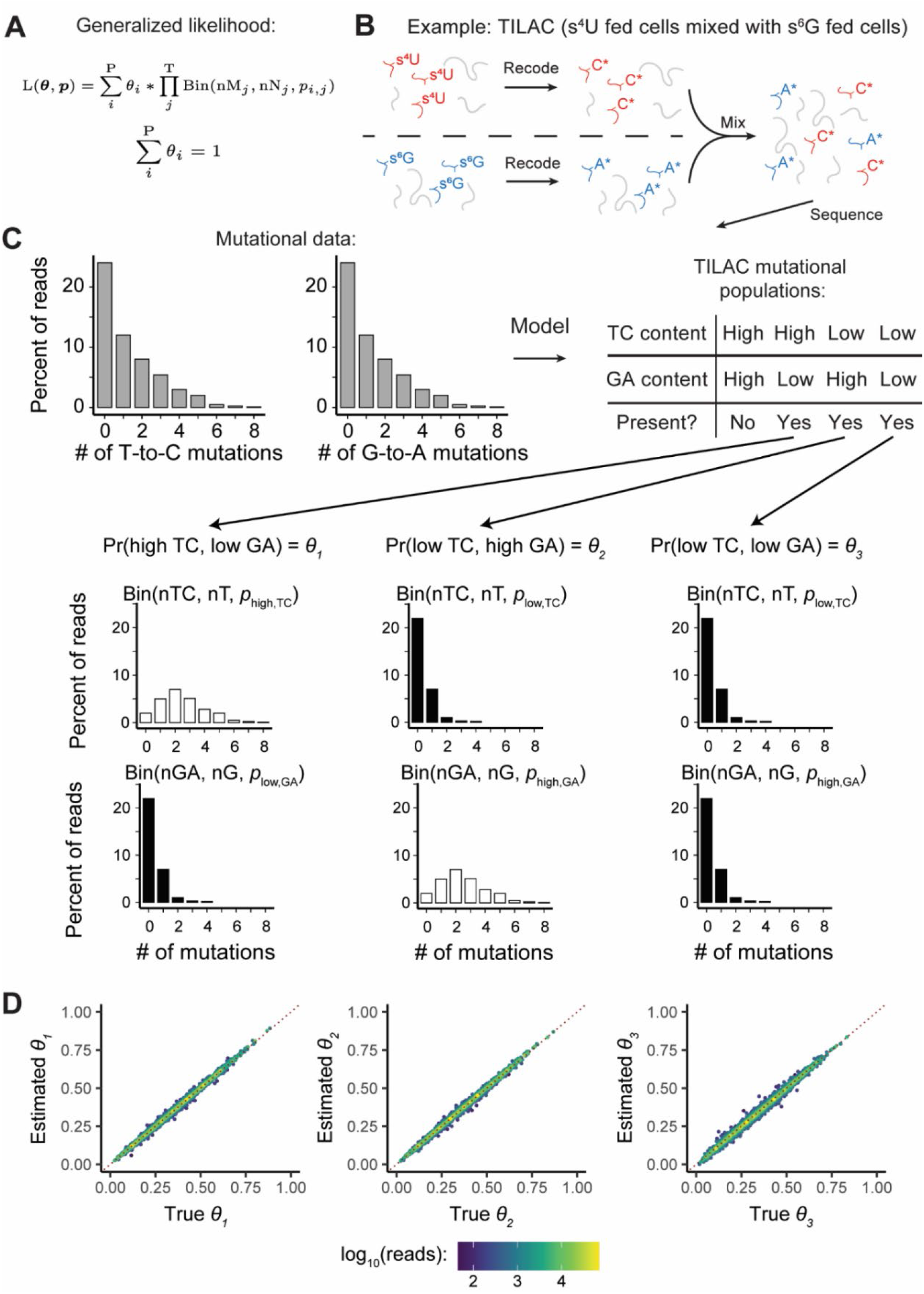
EZbakR generalizes NR-seq mixture modeling to support multi-label analyses. **A)** Generalized mixture model likelihood. P = number of distinct mutational populations (e.g., high T-to-C and low G-to-A mutation rate). T = number of mutation types being analyzed (e.g., T-to-C and G-to-A). nM = number of mutations of a particular type in a given read. nN = number of mutable nucleotides of a given type in a given read. **B)** Example of a dual-label NR-seq experimental method: TILAC. In this experiment, s^4^U fed cells are mixed with s^6^G fed cells. **C)** Schematic for how generalized mixture modeling works in the setting of TILAC. In TILAC, there are no dually labeled reads, so the high T-to-C and high G-to-A population does not exist (see mutational populations table). **D)** Analyses of simulated TILAC data. *θ*_1_ = fraction s^4^U labeled; *θ*_2_ = fraction s^6^G labeled; *θ*_3_ = fraction unlabeled. X-axis is simulated ground truth. Y-axis is estimated value. Red dotted line is perfectly accurate estimation.

GRAND-SLAM and bakR make several additional assumptions regarding the mutational content of sequencing reads. Most significantly, they both assume that all RNA synthesized in the presence of metabolic label had an equal probability of incorporating said label. This is codified in the assumption that the mutational distribution for labeled reads is well described as a binomial distribution with one probability of mutation parameter (*p*_labeled_) for all reads in a given sample. Recent work has shown that this assumption is violated by some transcript types. For example, mitochondrial RNAs have much lower labeled read mutation rates than other RNAs (McShane, et al., 2024). Because of this, we developed a strategy to allow each analyzed feature to have its own incorporation rate parameter. To make this strategy robust, we employed a hierarchical modeling approach, in which we first infer a conservative feature-specific *p*_labeled_ prior distribution, and use this to regularize the feature-specific estimates (Fig. 4A and Methods). This prevents extreme estimate variance due to low feature coverage limiting the precision of the feature-specific incorporation rate estimation. Analyses of simulated data confirmed that this strategy provides accurate estimates of the fraction labeled and feature-specific *p*_labeled_ (Fig. 4B). Analyzing a real TimeLapse-seq dataset revealed that this strategy is able to identify and account for the previously-reported low incorporation rate for transcripts encoded on the mitochondrial chromosome (Fig. 4C) (Ietswaart, et al., 2024; McShane, et al., 2024). While the hierarchical and non-hierarchical models provide similar fraction labeled estimates in most cases, the hierarchical model outputs far higher fraction labeled estimates for these mitochondrially encoded transcripts (Figure S2). This is due to a sample-wide *p*_labeled_ being an overestimate for these transcripts, effectively leading to the misclassification of many reads with moderate T-to-C mutation counts as unlabeled when using the non-hierarchical model (Figure 4C). The EZbakR suite thus improves the mixture modeling of NR-seq data.

**Figure 4:**
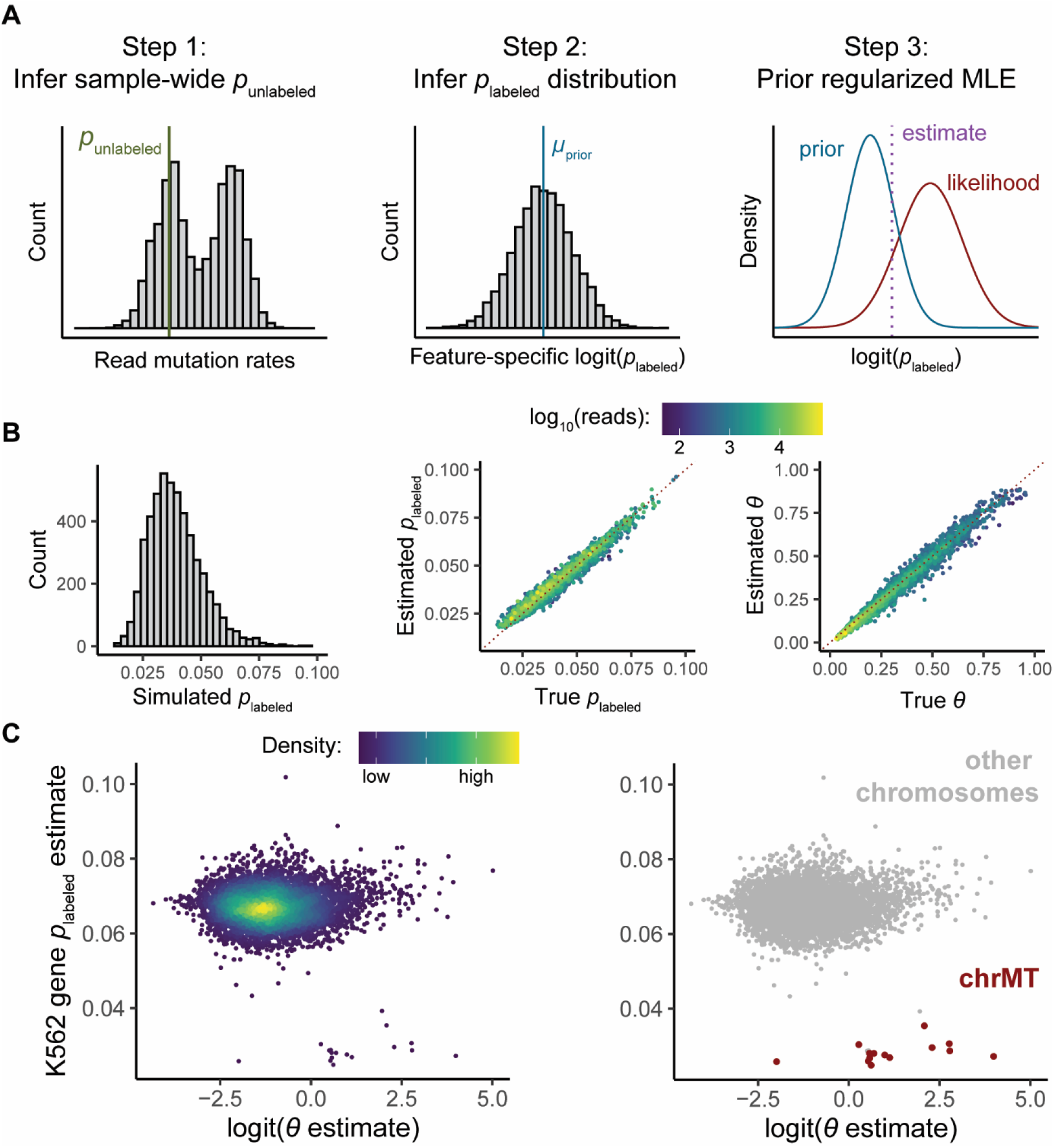
EZbakR’s hierarchical NR-seq mixture modeling accounts for *p*_labeled_ variation. **A)** Schematic of hierarchical modeling strategy to infer a *p*_labeled_ for each feature (i.e., feature-specific *p*_labeled_). Strategy is designed to strongly regularize feature-specific *p*_labeled_ estimates to reduce estimate variance. See Methods for details. **B)** Analyses of simulated data. Left: distribution of simulated feature-specific *p*_labeled_. Middle: assessment of feature-specific *p*_labeled_ estimate accuracy. Right: Assessment of fraction labeled (*θ*) estimate accuracy. In Middle and Right plots, red, dotted line represents perfect estimation. Points are colored by simulated read count. **C)** Estimated feature-specific *p*_labeled_ (Y-axis) as a function of the estimated fraction labeled (on a logit-scale; X-axis) from analysis of TimeLapse-seq data from K562 cells (Ietswaart et al., 2024). Left: points colored by density. Right: points colored by whether RNA originated from the mitochondrial chromosome (chrMT).

### Generalized linear dynamical systems modeling

Once the labeled read abundance for a given feature has been rigorously estimated, the next step of most NR-seq analyses is the inference of kinetic parameters. In a standard NR-seq analysis, this means using the exonic fraction labeled and normalized read count to infer synthesis and degradation rate constants. Implicitly, this means assuming a model of the RNA dynamical system where mature mRNA is synthesized at a rate k_syn_ and degraded with a rate constant k_deg_ (Figure 5A). Recently, several studies have combined subcellular fractionation with NR-seq to investigate the rate of flow between subcellular compartments, as well as to probe the dynamics of RNA in each compartment (Ietswaart, et al., 2024; Müller, et al., 2024; Steinbrecht, et al., 2024). In this case, the data needs to be fit to a more complicated model. To date, this has required in-house, specialized solutions for a given fractionation scheme and assumed dynamical system. A strategy to generalize this approach to support any experimental design and model of RNA population dynamics would greatly facilitate analyses of these data.

**Figure 5:**
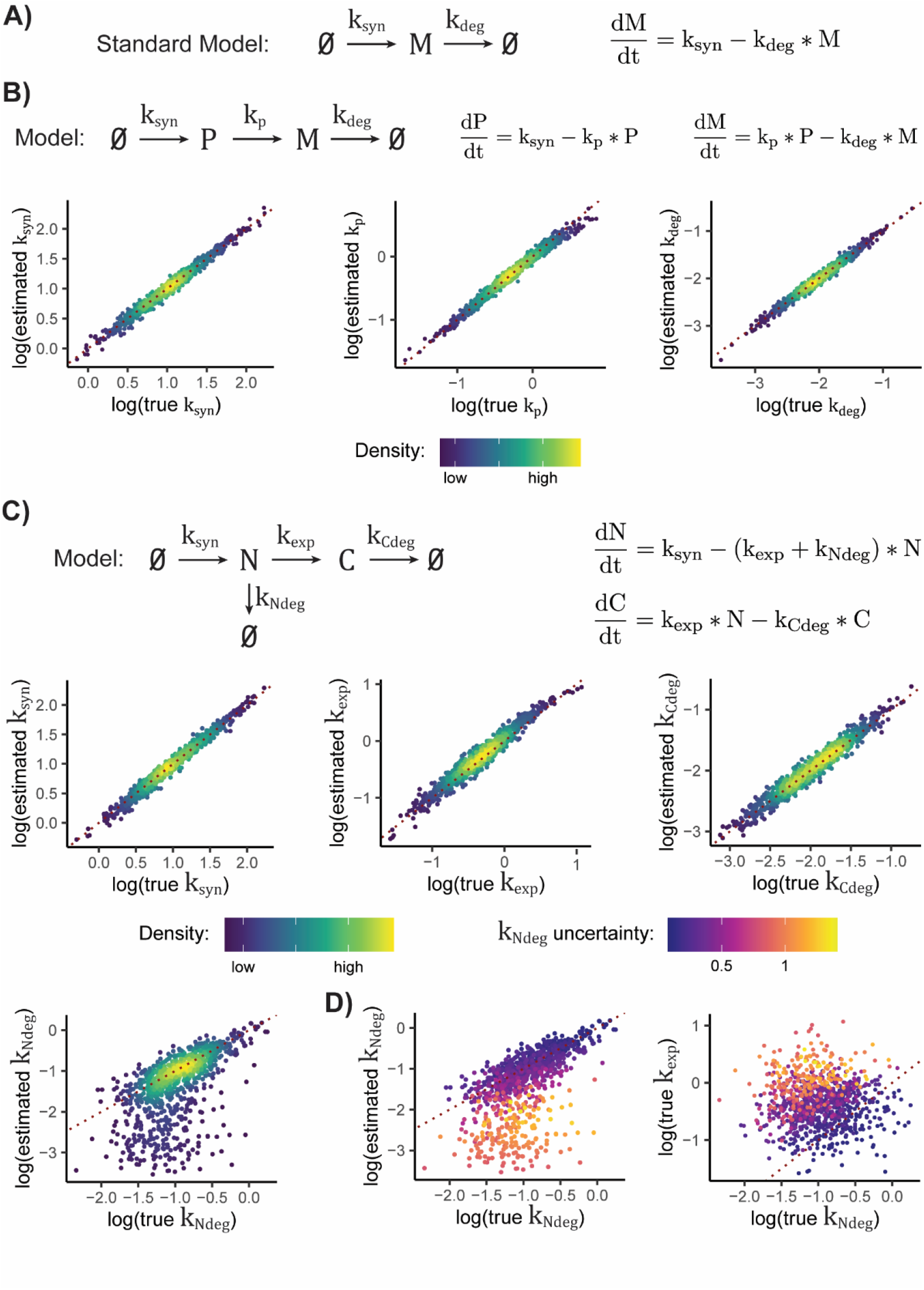
EZbakR generalizes kinetic modeling of NR-seq data. **A)** Model assumed when performing standard analysis of mature mRNA synthesis and degradation. **B)** Analysis of simulated data for model of pre-mRNA maturation. P = premature mRNA; M = mature mRNA. Scatter plots show comparison of true simulated parameter values to those estimated by EZbakR, for all three kinetic parameters in said model. Red dotted line represents perfect estimation. **C)** Analysis of simulated data for model of nuclear-to-cytoplasmic trafficking of RNA. N = nuclear RNA. C = cytoplasmic RNA. Red dotted line represents perfect estimation. **D)** Left: Nuclear degradation rate constant accuracy scatterplot from **C**, colored by model’s uncertainty in rate constant estimate. Right: comparison of the true nuclear degradation and export rate constants, colored by model’s uncertainty in nuclear degradation rate constant. Red dotted line represents equal nuclear degradation and export kinetics. Estimating k_Ndeg_ is expected to get harder the further points are from this line, for reasons discussed in Supplemental Methods.

Towards that end, EZbakR allows users to specify and estimate parameters of any identifiable, linear dynamical systems model of their choice. All such models, depicted as graphs in Figure 5, can be described using a system of ordinary differential equations (ODEs) with analytic solutions (see Methods and Supplemental Methods for details). EZbakR infers the analytic solution for the dynamics of all modeled RNA species and uses this to perform maximum likelihood estimation of the kinetic parameters in the specified model. By combining generalized feature assignment in fastq2EZbakR with generalized linear dynamical systems modeling, EZbakR is able to model premature mRNA processing dynamics. Analyses of simulated data confirm that this parameter estimation procedure is accurate (Fig. 5B). Despite the low absolute abundance of pre-mRNA, introns are typically much longer than exons, meaning that a significant portion of mapped reads in standard total RNA libraries originate from pre-mRNA (Figure S3). EZbakR allows users to make full use of all of these reads. Analyses of a real NR-seq dataset with this approach revealed that pre-mRNA processing kinetics are typically much faster than mature mRNA degradation kinetics, which is a conclusion well corroborated by previous work (Figure S3) (De Pretis, et al., 2015; Pai, et al., 2017; Shine, et al., 2024; Wachutka, et al., 2019).

EZbakR also supports analyses of RNA flow between subcellular compartments (Figure 5C) from experiments such as those described in Ietswaart et al., 2024. This requires integrating information across multiple independent samples from distinct RNA populations. Thus, using the read counts from these samples to inform kinetic parameter estimates would require a normalization strategy accounting for the differences in absolute molecular abundances across samples. While this is generally difficult, we drew inspiration from recent work and developed a normalization strategy that, with a sufficient set of samples, permits rigorous normalization (Steinbrecht, et al., 2024). This strategy relies on having samples from each individual fraction, as well as a necessary set of combinations of fractions (e.g., nuclear, cytoplasmic, and whole cell RNA). See Methods and Supplemental Methods for details, but in short, scale factors can be derived from modeling the total fraction of reads that are labeled in the combination of fractions (e.g., whole cell data) as a weighted sum of the same quantity in each individual fraction (e.g., nuclear and cytoplasmic fractions). Analyses of simulated nuclear and cytoplasmic fractionation data confirm that this strategy is accurate (Figure 5C). In addition, using the normalized read counts improves the accuracy of kinetic parameter estimates in simple models, while also permitting investigation of more complicated models (Figure S4 and S5). One challenge of fitting more complex models is that while all parameters may be theoretically identifiable, they may not be practically identifiable, e.g., due to the limited impact that the precise value a given parameter has on measured quantities (Figure 5D; also see Supplemental Methods for further discussion). EZbakR thus provides uncertainty quantification that can flag instances of practical unidentifiability, allowing users to avoid overinterpreting low confidence estimates (Figure 5D). Applying this strategy to a recent subcellular fractionation NR-seq dataset allowed us to explore the prevalence of nuclear degradation in a human cell line (Figure S6). EZbakR corroborated many suspected genes producing a significant number of RNAs degraded in the nucleus (recently termed predicted to undergo nuclear degradation, or PUNDs), while also expanding the list of putative PUNDs (Supplemental Table 1) (Ietswaart, et al., 2024). The EZbakR suite thus generalizes the kinetic modeling of NR-seq data.

### Generalized comparative analyses

Often, important biological conclusions are drawn not from kinetic parameter estimates in a given experimental condition, but from comparative analyses of these estimates across different biological conditions. bakR introduced a novel hierarchical model that significantly increased the power of comparative analyses of synthesis and degradation kinetics (Vock and Simon, 2023). We have implemented this model in EZbakR, and it is compatible with all of the kinetic parameter inference strategies in EZbakR. While bakR is only able to perform comparisons of multiple experimental conditions to a single reference condition, EZbakR implements a design matrix-specified generalized linear model that greatly increases the flexibility of its comparative analyses (Figure 6A). In addition to providing greater model flexibility, EZbakR also improves upon the computational performance of bakR. bakR includes three implementations of the same model, which differ in statistical rigor and computational intensiveness (Vock and Simon, 2023). It includes a fast but conservative implementation (termed the MLE implementation) as well as more highly powered implementations that suffer from much longer runtimes (e.g., the MCMC implementation). In EZbakR, we have improved upon the efficient implementation in bakR so as to achieve statistical performance on-par with the more computationally intensive strategies in bakR, while not sacrificing efficiency (Figure 6B-E). The EZbakR suite thus generalizes performing well-powered comparisons of RNA kinetic parameters with NR-seq data.

**Figure 6:**
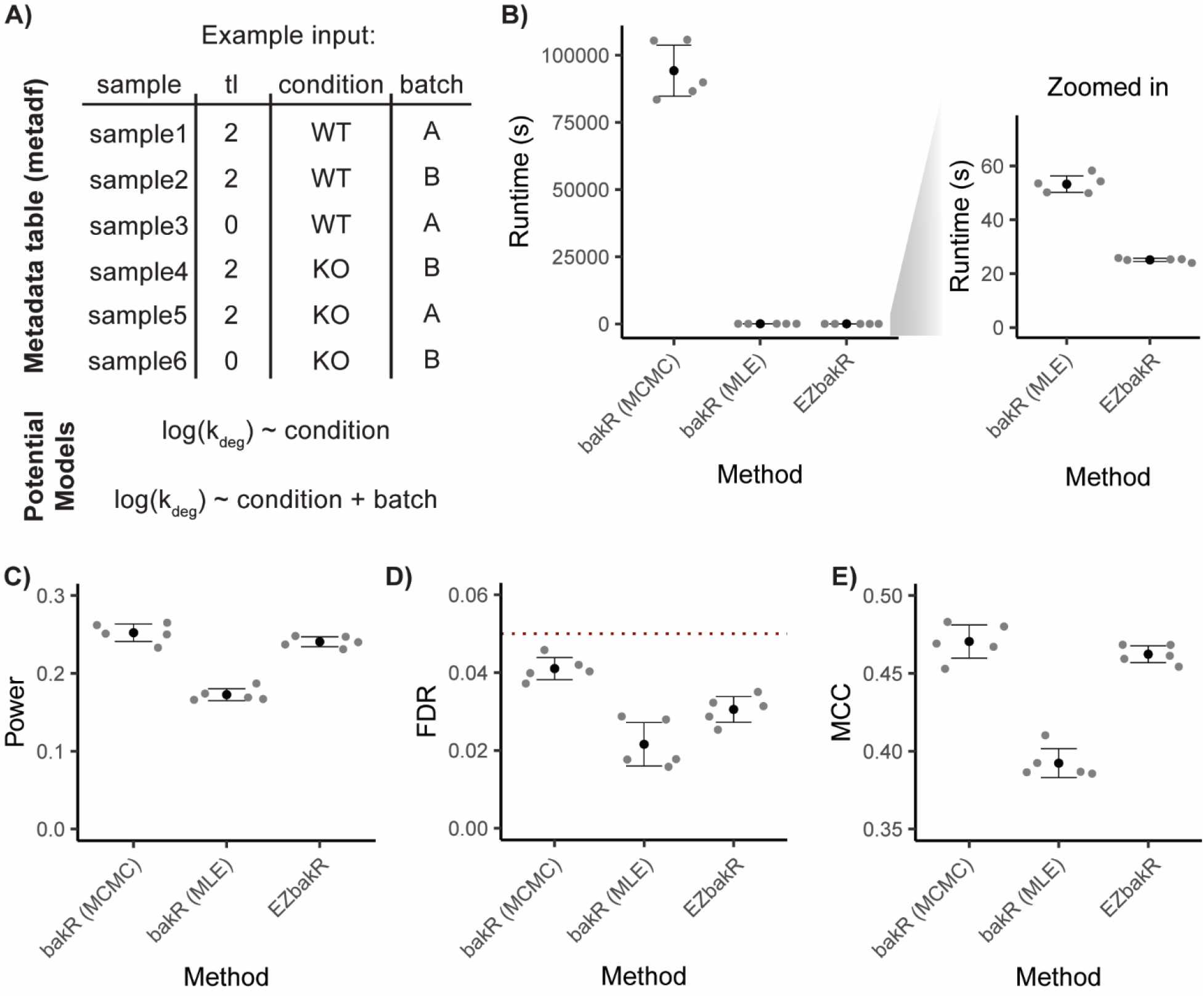
EZbakR improves and generalizes performing comparative analyses with NR-seq. **A)** Input to linear model of kinetic parameters in EZbakR. Includes metadata for each sample analyzed and a model relating a given kinetic parameter to factors included in metadata. Any identifiable model can be specified and fit. This approach allows for simple multi-condition comparisons (top potential model) or more complicated analysis designs (e.g., batch effect modeling; bottom potential model) **B)** Comparison of runtimes between two bakR implementations (Markov Chain Monte Carlo (MCMC) and Maximum Likelihood Estimation (MLE)) and EZbakR. **C-E)** Analysis of simulated data originally presented in Vock and Simon 2022. **C)** Comparison of statistical power (number of true positives / number simulated positives) between bakR implementations and EZbakR. **D)** Comparison of false discovery rates (FDRs; number of false positives / number of positives) between bakR implementations and EZbakR. **E)** Comparison of Matthew’s correlation coefficients (MCC) between bakR implementations and EZbakR.

## Discussion

To generalize and improve multiple aspects of NR-seq analyses we have developed the EZbakR suite, which includes a Snakemake pipeline (fastq2EZbakR) and an R package (EZbakR) (Figure 1). Unlike existing tools, the EZbakR suite facilitates analyses of a wide array of genomic features (Figure 2), implements strategies for multi-label analyses (Figure 3), offers improved single-label modeling (Figure 4), can fit any identifiable linear dynamical systems model to NR-seq data (Figures 5), and implements an efficient and flexible generalized linear model-based kinetic parameter comparison strategy (Figure 6). The EZbakR suite thus promises to greatly facilitate deriving novel insights from NR-seq data.

The EZbakR suite will be useful to perform in-depth analyses of new NR-seq datasets, and also has the potential to uncover biological signal initially missed in existing NR-seq datasets. The limitations of current NR-seq bioinformatic tools have forced most analyses of total RNA NR-seq datasets to focus solely on the dynamics of mature mRNA. The EZbakR suite’s generalized feature assignment combined with its generalized linear dynamical systems modeling opens the door for investigating the kinetics of RNA processing in unprecedented detail. In addition, by performing analyses at the levels of splice junctions, exonic bins, or transcript equivalence classes, the EZbakR suite has the potential to identify isoform-level effects missed by standard gene-wide aggregate analyses. In doing so, the EZbakR suite will allow researchers to make full use of their NR-seq data.

The EZbakR suite is designed to flexibly support analyses of the full universe of existing and future NR-seq extensions. From dynamical systems modeling of flow between subcellular compartments to analyses of multi-label NR-seq datasets, the EZbakR suite directly facilitates extension-specific analyses that no other tool supports. It is difficult for a single tool to perform all possible downstream analyses. For example, RNA velocity analyses of single cell NR-seq datasets both remain beyond the scope of the EZbakR suite and are supported by a number of existing specialized tools (Maizels, et al., 2024; Qiu, et al., 2022; Weiler, et al., 2024). That said, all NR-seq analyses must start with the quantification of labeled RNA species. By improving models to perform these analyses, and providing convenient access to the data required to fit such models, the EZbakR suite promises to improve analyses of all NR-seq datasets, regardless of their particular idiosyncrasies (e.g., unique data preprocessing requirements). In addition, the modular design of the EZbakR suite will facilitate future development and extension of its functionalities. We anticipate that the EZbakR suite will provide a unified bioinformatic platform to support the ever-expanding NR-seq ecosystem.

## Methods

### fastq2EZbakR

fastq2EZbakR is a Snakemake pipeline that can accept either fastq files or aligned bam files (the latter making fastq2EZbakR compatible with specialized alignment strategies not directly implemented in our pipeline (Berg, et al., 2024)) as input (Figure 2B) (Köster and Rahmann, 2012). It produces as output a file we term a <u>c</u>ounts <u>B</u>inomial (cB) file, which contains information about the mutation content of reads and the features they were assigned to (see Figure 1, highlight 2 for simplified schematic). If fastq files are provided as input, adapters are first trimmed with fastp and reads are aligned with either STAR or HISAT2 (user choice, STAR by default given recent benchmarks showing it performs similarly to specialized 3-base alignment strategies) (Chen, et al., 2018; Dobin, et al., 2013; Kim, et al., 2019; Popitsch, et al., 2024; Zhang, et al., 2021). In all cases, unmapped reads and non-primary alignments are filtered from the resulting or provided bam file with SAMtools (Bonfield, et al., 2021). Mutations are counted with a custom Python script, and feature assignment is performed as described in the Supplemental Methods under *Generalized feature assignment in fastq2EZbakR* (all strategies are schematized in Figure 2A), using a combination of featureCounts and custom scripting (Liao, et al., 2014). The mutation counts and feature assignment tables are merged and combined for all samples to give the final cB file. Documentation detailing how to run fastq2EZbakR can be found here: https://fastq2ezbakr.readthedocs.io/en/latest/

### Generalized mixture modeling in EZbakR

Mutational mixture modeling in EZbakR is performed in two steps:

1. Mutation rates in labeled and unlabeled reads (*p*_labeled_ and *p*_unlabeled_) for all types of mutations being modeled are estimated. This is done by fitting a two-component mixture model for each mutation type to all reads from a given sample. Fitting is done for each mutation type independently at this stage. If users have control samples lacking any labeling, they can optionally choose to use a single *p*_unlabeled_ derived from these samples. This can improve the stability of *p*_labeled_ estimates in some settings, like when incorporation rates are low.
2. The generalized mixture model parameters are estimated via the method of prior-penalized maximum likelihood, similar to as previously described for bakR (Vock and Simon, 2023).

See Supplemental Methods for further details and mathematical formalization (formalization is also succinctly depicted in Figure 3A).

### Hierarchical mixture model in EZbakR

Hierarchical mixture modeling in EZbakR is currently only compatible with standard two-component mixture modeling (Figure 4A). This strategy starts with the same step 1 as in the generalized mixture modeling described above. After global estimates for *p*_labeled_ and *p*_unlabeled_ are obtained, the following iterative strategy is used to perform maximum likelihood estimation with feature-specific *p*_labeled_ values:

1. A two-component mixture model with feature-specific *p*_labeled_’s is fit to all features with more than a certain number of reads (300 by default) in each sample. Low coverage features are not considered in this step as the limited number of reads for such features precludes stable estimation of a feature-specific *p*_labeled_.
2. A strongly regularizing logit(*p*_labeled_) prior is estimated for each sample from the distribution of feature-specific *p*_labeled_’s estimated in the previous step. More specifically, a normal distribution prior is inferred with mean equal to the estimated sample-wide logit(*p*_labeled_) and standard deviation equal to the standard deviation of feature-specific logit(*p*_labeled_) estimates minus the average feature-specific logit(*p*_labeled_) uncertainty. If this difference is less than 0, a user-defined small standard deviation (0.01 by default) is imputed. If this difference is greater than a user-defined maximum (0.15 by default), the maximum is imputed. Extreme estimates (i.e., those at the set parameter bounds of between -9 and 0 on the logit scale) are filtered out before calculating the mean and standard deviation of the distribution. If multiple labeled samples are being modeled, the minimum sample-specific prior standard deviation is used for all samples so as to avoid under-conservativeness that could yield highly unstable estimates in the next step.
3. A two-component mixture model with feature-specific *p*_labeled_’s is fit to all features with the strongly regularizing logit(*p*_labeled_) prior obtained in step 2.

### Generalized linear dynamical systems modeling in EZbakR

Generalized linear dynamical systems modeling in EZbakR is performed with the EZDynamics() function. It can take as input either sample-specific fraction labeled estimates (output of EZbakR’s EstimateFractions() function) or condition-wide averages obtained from the EZbakR generalized linear model fit (AverageAndRegularize() function). Both are compatible with modeling of pre-mRNA dynamics, but modeling of RNA flow between subcellular compartments is only compatible with the latter. This is because modeling of subcellular compartment flow requires integrating across multiple independent samples (i.e., data from different subcellular fractions), so there is no way to estimate all kinetic parameters of such models for individual samples. The input to EZDynamics() is a matrix representation of graphical models akin to those in Figure 5. Documentation regarding model specification can be found here: https://isaacvock.github.io/EZbakR/articles/EZDynamics.html. Other input includes information about how the measured features relate to the modeled features. For example, if fitting the subcellular fractionation model shown in Figure 5C, you will likely have whole cell data. The RNA from this sample corresponds to the sum of N (nuclear RNA) and C (cytoplasmic RNA), and this fact must be conveyed to EZDynamics(). EZDynamics() then uses the method of maximum likelihood to estimate kinetic parameters of your model, and uncertainties are calculated as the square root of the diagonal elements of the inverse Hessian matrix (Fonseca, et al., 2005). This is done by modeling both the fraction labeled and, if rigorous normalization is possible, the read counts for a given feature. See Supplemental Methods for details.

Estimating scale factors for multi-compartment modeling can be done in one of three ways:

1. Spike-in normalization. EZbakR allows users to input spike-in derived scale factors.
2. Fraction labeled modeling. The fraction labeled estimates (which are internally normalized) can be used to estimate all but the RNA synthesis parameter, if such an estimation strategy is identifiable. Scale factors can then be inferred from the set of non-synthesis parameter estimates, by identifying the factors by which absolute RNA abundances are expected to differ across the compartments (since the synthesis rate is the same for all RNA species from a given gene). A downside of this strategy is that it limits the complexity of identifiable models, and typically yields lower confidence parameter estimates (Figure S4, top row).
3. Fraction labeled mixing model. If users have data for individual subcellular compartments as well as data for a sufficient number of combinations of these compartments, then scale factors can be estimated from the differences in overall fraction of reads that are labeled in each compartment or combination of compartments. For example, if users have nuclear fraction, cytoplasmic fraction, and whole cell data, then normalization factors can be estimated by solving a linear system of equations and using the inferred ratio of absolute molecular abundances to derive scale factors for each compartment. See Supplemental Methods for details and mathematical formalization.

Option 3 was used in analyses of subcellular fractionation data in Figures 5, S4, S5, and S6.

### Generalized linear modeling in EZbakR

EZbakR’s AverageAndRegularize() function is used to fit a generalized linear model of a user’s fraction labeled or kinetic parameter estimate data. Parameter standard error estimates are then regularized using a hierarchical modeling strategy similar to that introduced in bakR (Vock and Simon 2022). See Supplemental methods for a discussion of the improvements made to this regularization scheme in EZbakR. The EZbakR input data object (an EZbakRData object) includes a so-called metadata data frame (metadf), which can contain any number of factors describing elements of each sample (see example in Figure 6A). These factors can be included in a formula object passed to AverageAndRegularize(), which will specify the linear model to fit to one’s data. A unique aspect of generalized linear modeling in EZbakR is that it can allow for heteroskedasticity through a number of different ways. One, if a model is such that each parameter is estimated from a non-overlapping set of samples, then standard deviations are estimated for each of these sample sets independently or two, users can specify a set of factors by which to group samples, and standard deviations are estimated in these sample sets and these standard deviations are used to estimate parameter standard errors. See Supplemental Methods for details.

### Generating simulated data

Simulated data used in this study were generated with simulation functions implemented in EZbakR. Multi-label data was simulated with the SimulateMultiLabel() function in EZbakR. Data with feature-to-feature *p*_labeled_ variation was simulated with the SimulateOneRep() function in EZbakR. Data for all dynamical systems models tested in Figure 4 were simulated with the SimulateDynamics() function in EZbakR. See documentation and Supplemental methods for details regarding how these functions work and parameters used. Simulated data for analyses in Figure 6 came from the previously published bakR benchmarks (Vock and Simon, 2023).

## Data availability

Fastq2EZbakR and EZbakR are both freely available on Github:

fastq2EZbakR: https://github.com/isaacvock/fastq2EZbakR/blob/main/README.md

EZbakR: https://github.com/isaacvock/EZbakR

Scripts to reproduce all figures and supplemental figures are available on Github, at: https://github.com/isaacvock/EZbakRsuite_paper_code

Processed data necessary to reproduce all figures is available on Zenodo (DOI: 10.5281/zenodo.13929898), as well as supplemental tables of:

- All kinetic parameter estimates for analysis of total-cytoplasmic-nuclear NR-seq data (Figure S6).
- All kinetic parameter estimates for analysis of premature mRNA and mature mRNA dynamics (Figure S3).

## Acknowledgements

This work was funded by National Institutes of Health (NIH) grants R01GM137117, T32GM67543-19, and NHLBI ZIAAHL006158. This work was also supported by the Intramural Research Program, National Heart, Lung, and Blood Institute, National Institutes of Health, and utilized the computational resources of the NIH HPC Biowulf cluster (http://hpc.nih.gov).

The authors thank members of the Simon (namely Andreas Pintado-Urbanc and Michelle Moon), Pai (namely Jesse Lehman and Athma Pai), and Churchman (namely Robert Iestwaart, Erik McShane, Mary Couvillion, and Stirling Churchman) labs for useful discussions during the building and testing of EZbakR. We also thank the Churchman lab for early access to their subcellular TimeLapse-seq dataset. Finally, we want to thank all members of the Simon lab for helpful comments on this manuscript.

## Author contributions

IWV developed the EZbakR suite, wrote the manuscript, and performed all analyses. IWV and MDS conceived the project. IWV, JWM, AZ, MM, and JRH conceived the various feature assignment strategies. JWM and AZ tested the EZbakR suite and suggested a number of analyses. MDS and JRH provided funding support. All authors helped edit the manuscript.

