## Supplemental Figures and Methods for "Expanding and improving analyses of nucleotide recoding RNA-seq experiments with the EZbakR suite"

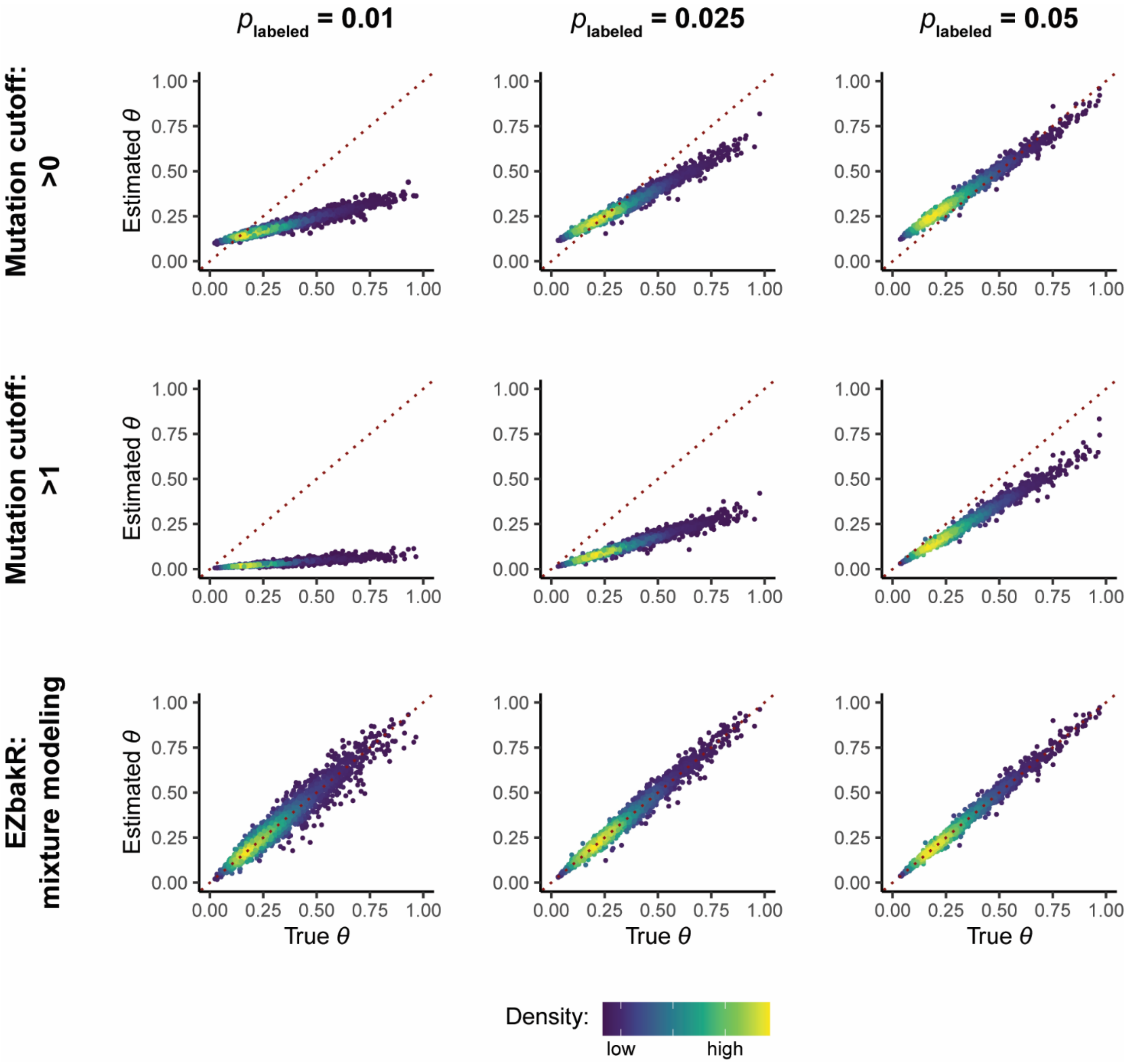


Figure S1: Simple mutation cutoff analysis strategies are less accurate than mixture modeling. Comparison of accuracy of 3 analysis strategies on 3 datasets. Each column represents analysis for a particular dataset. Datasets differ in their simulated $p_{\text{labeled}}$ (1%, 2.5%, and 5% mutation rates). Each row represents a particular analysis strategy. **Top row:** labeled reads are defined as those with at least 1 (> 0) mutation. This is the strategy implemented in SLAMDUNK by default. **Middle row:** labeled reads are defined as those with at least 2 (> 1) mutations. Represents another commonly used NR-seq analysis cutoff. **Bottom row:** EZbakR analysis with two-component mixture modeling. Points are colored by density in all plots. $\theta$ = fraction of reads from labeled RNA. Red dotted lines represent perfect estimation.

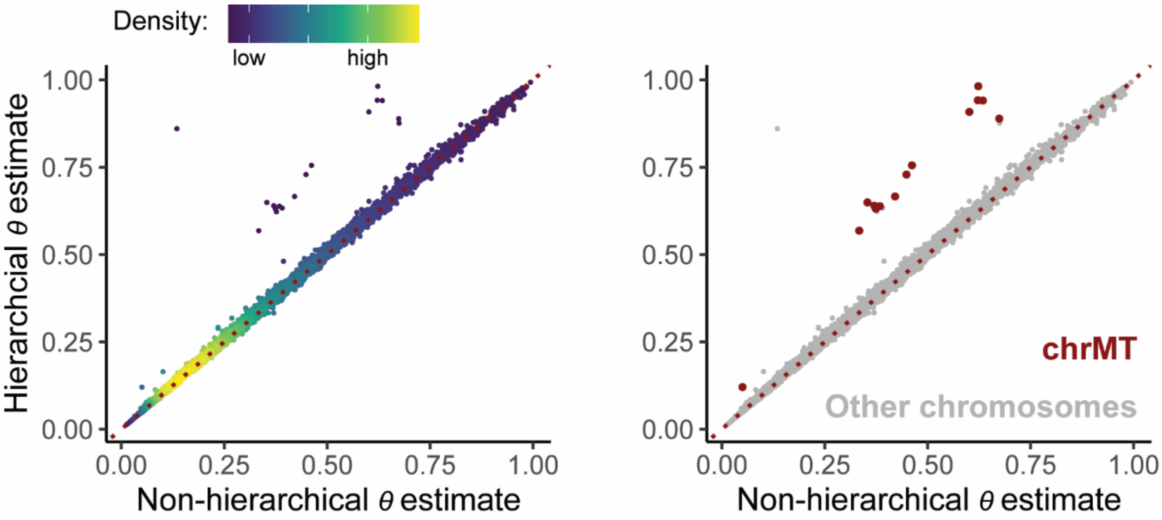


Figure S2: Comparison of fraction labeled ($\theta$) estimates with and without the $p_{\text{labeled}}$ hierarchical model introduced in Figure 3. X-axis represents estimates from standard, non-hierarchical $p_{\text{labeled}}$ model (single $p_{\text{labeled}}$ used for all genes in a given sample). Y-axis represents estimates from novel, hierarchical $p_{\text{labeled}}$ model. **Left:** Points colored by density. **Right:** points colored by whether or not the gene is on the mitochondrial chromosome (chrMT). Analyses are of a real total RNA TimeLapse-seq dataset (Ietswaart, et al., 2024).

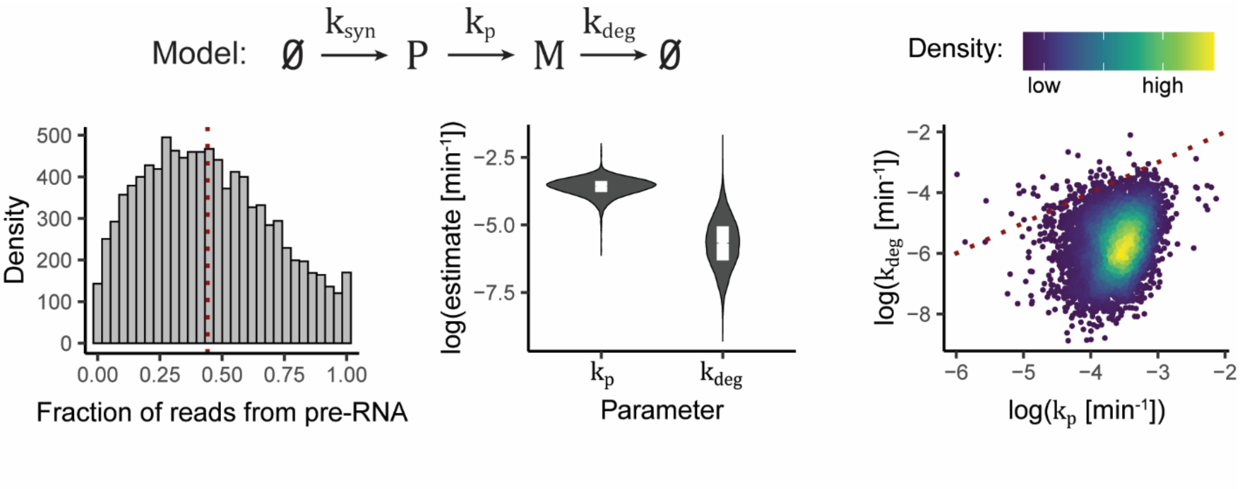


Figure S3: EZbakR can fit model of pre-mRNA dynamics to real data. Analysis of whole cell TimeLapse-seq data, fitting model presented in Figure 4A and at the top left of this figure. **Left:** Fraction of reads from each gene that are likely derived from pre-mRNA (are assigned to a gene but don't exclusively overlap with exonic regions of said gene). Fraction pre-mRNA's of 1 typically represent annotation discrepancies. These are not analyzed as EZbakR requires representation of pre-mRNA and mature RNA reads to estimate all parameters in the model. **Middle:** Distributions of processing and mature RNA degradation rate constants. **Right:** Comparison of gene-specific processing and mature RNA degradation rate constant estimates. Points colored by density, with brighter colors representing higher density.

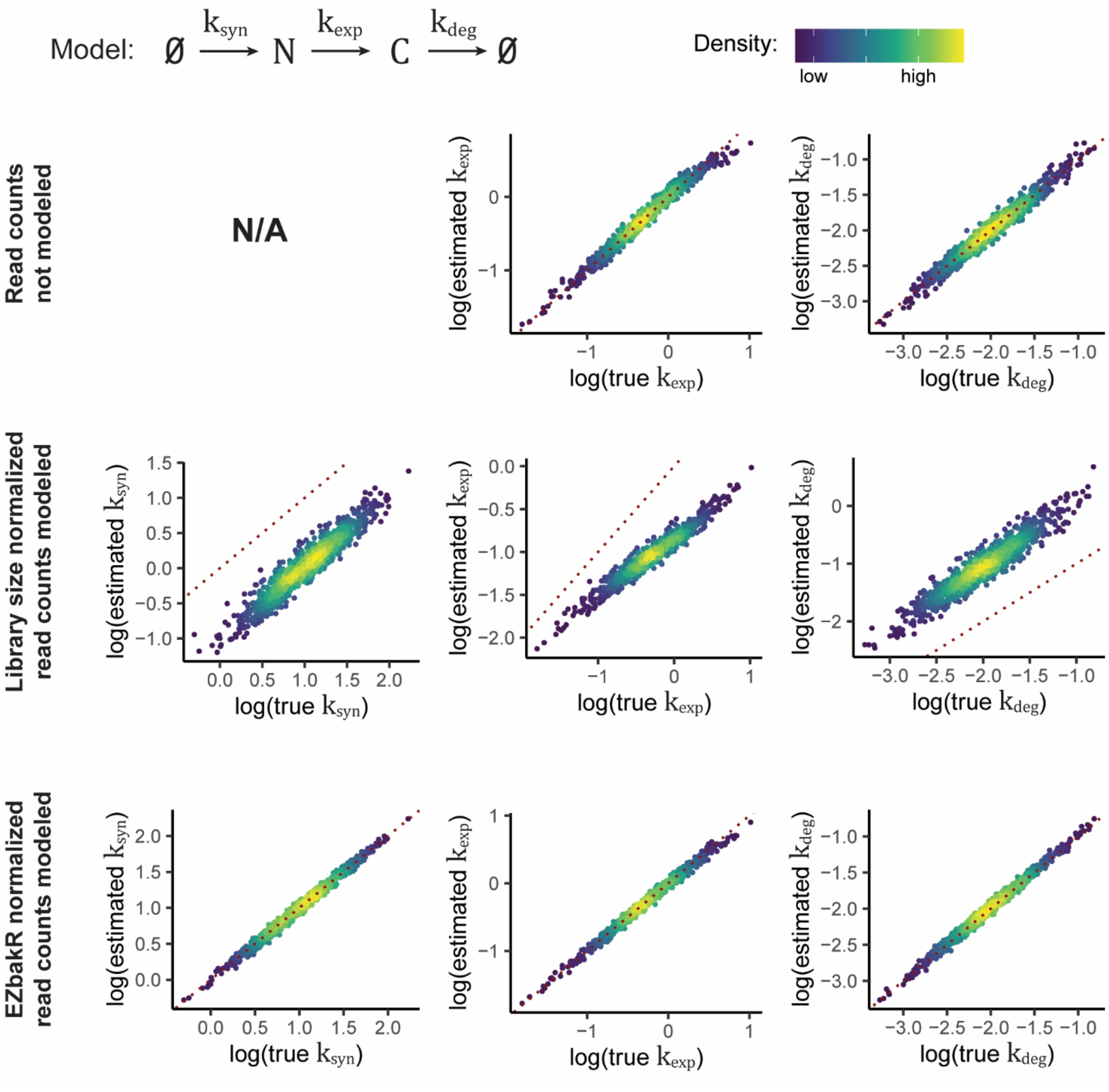


Figure S4: Proper read count normalization improves analyses of subcellular RNA flow. The model presented at the top (N = nuclear RNA, C = cytoplasmic RNA) was fit with three strategies, and the accuracy of kinetic parameter estimates were assessed in all three cases. **Top:** Read counts were not modeled. This precludes estimation of ksyn. **Middle:** A naïve library size normalization scale factor was applied to read counts, and these normalized read counts were used by EZbakR to estimate kinetic parameters. **Bottom:** Normalization scale factor was estimated using naïve strategy implemented in EZbakR and discussed in the Methods and Supplemental Methods, and these normalized read counts were used by EZbakR to estimate kinetic parameters. In all cases, points are colored by density, and red dotted line represents perfect estimation.

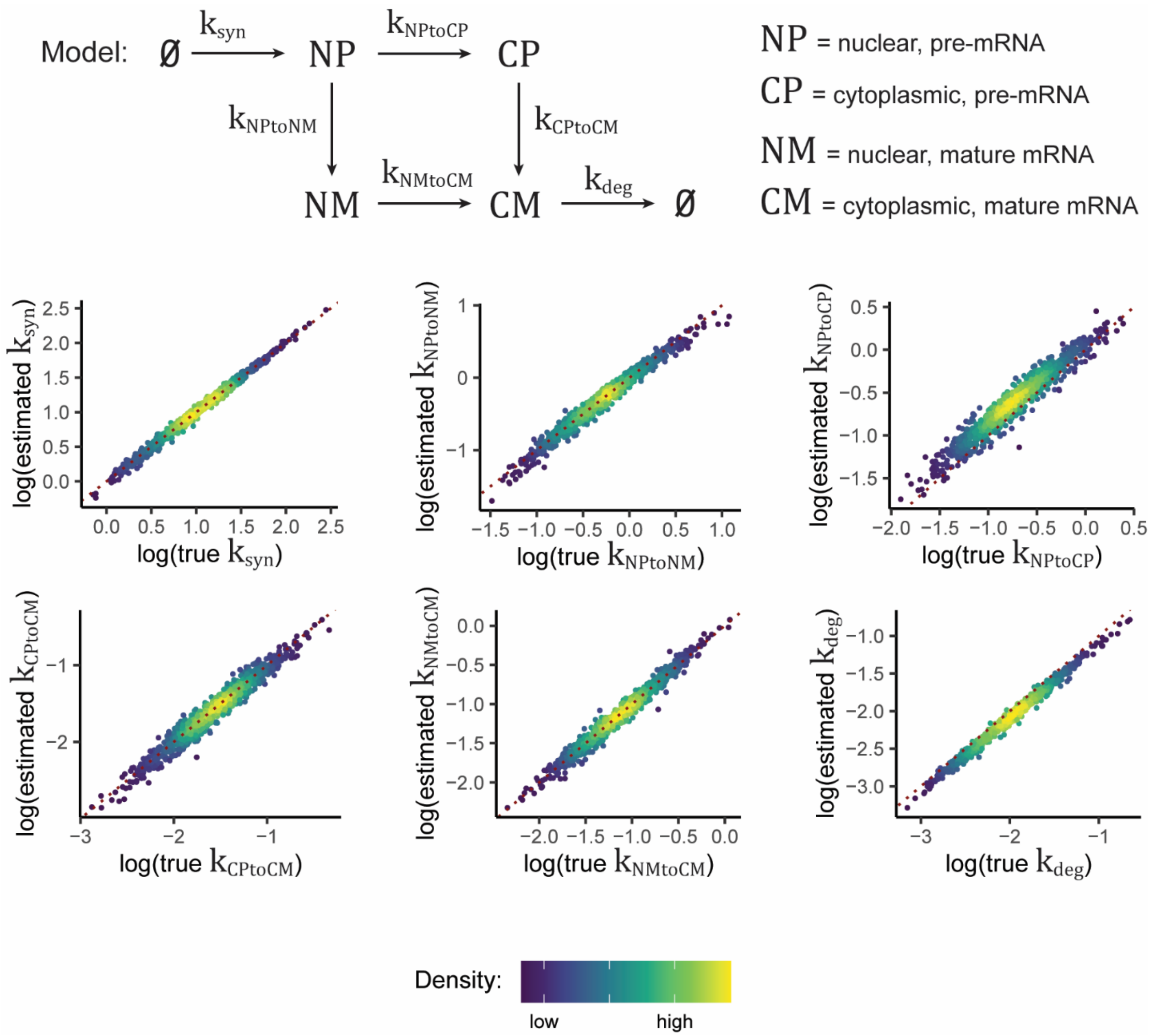


Figure S5: EZbakR can fit more complex models of premature and mature mRNA transport. Model at top of figure was simulated, and fit to said simulated data. The six scatterplots assess the accuracy of estimates for all six kinetic parameters in this model. In all cases, points are colored by density, and the red dotted line represents perfect estimation.

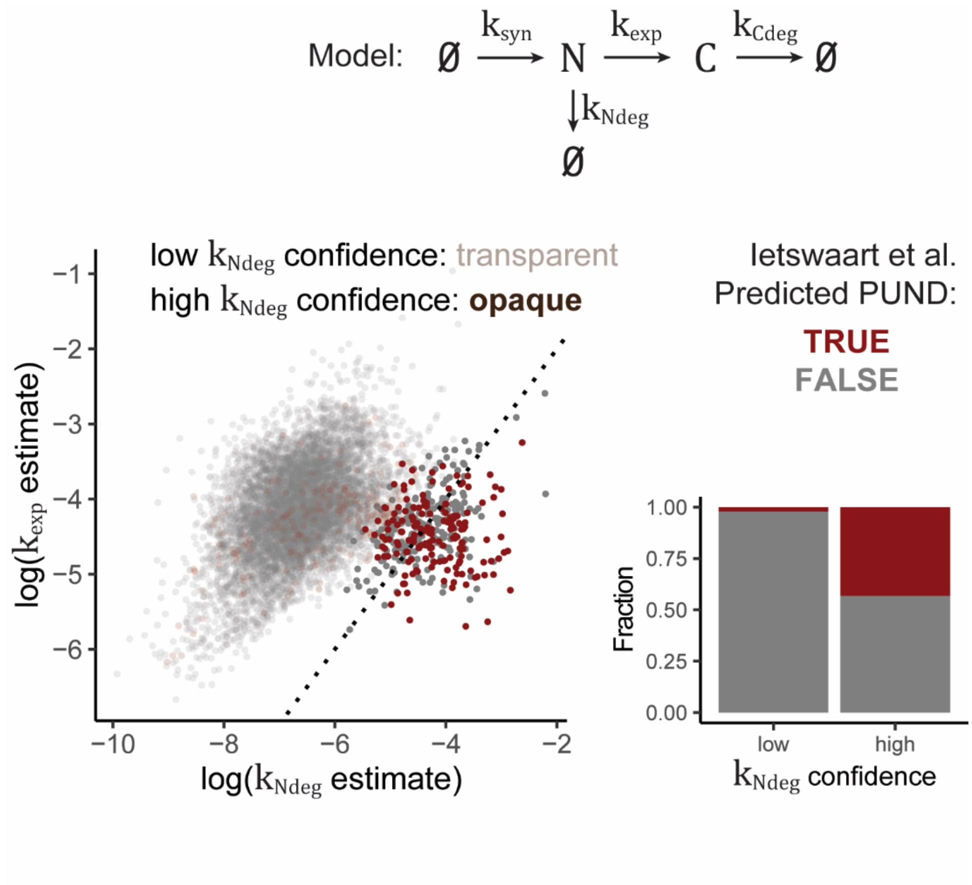


Figure S6: EZbakR identifies high confidence instances of significant nuclear degradation. Kinetic model at top of figure was fit to nuclear/cytoplasmic/whole cell data from Ietswaart et al., 2024. Estimated nuclear degradation rate constants were compared to nuclear export rate constants, with points colored via a cutoff in the $k_{Ndeg}$ uncertainty (bottom left scatter plot; > 0.2 is transparent). List of high confidence PUNDs (those with high confidence $k_{Ndeg}$ estimates, as these represent instances where $k_{Ndeg}$ is similar order of magnitude to $k_{exp}$) was compared with those identified previously (bottom right bar plot).

**Supplemental Methods**

*Generalized feature assignment in fastq2EZbakR*

As schematized in Figure 2A, fastq2EZbakR can assign reads to any combination of the following features: genes (anywhere), genes (exonic only), exonic bins, exon-exon junctions, and transcript equivalence classes. In all cases, feature assignment is performed with either featureCounts, custom scripting, or a combination of the two (Liao, et al., 2014). Below, describe these assignment strategies in detail:

1. Genes (anywhere): featureCounts is used to assign reads to anywhere in a gene. The following parameters are used by default in the call to featureCounts: `-R CORE -g gene_id -t transcript -M`. If paired-end reads are provided, the following is automatically added to the featureCounts parameters: `-p –countReadPairs`. The output CORE file is used in all downstream steps of the pipeline. This strategy will assign a read to a gene if it overlaps any portion of said gene. In this case, reads overlapping multiple annotated genes will go unassigned (value in relevant cB column of "__no_feature").
2. Genes (exonic only): featureCounts is used to assign reads to exclusively exonic regions of a gene. This means that reads overlapping any non-exonic region should be flagged as unassigned. The parameters used for this by default are: `--nonOverlap 0 -R CORE -g gene_id -M`. If paired-end reads are provided, the following is automatically added to the featureCounts parameters: `-p –countReadPairs`. The output

CORE file is used in all downstream steps of the pipeline. With this strategy, reads overlapping a region that is intronic in all annotated isoforms will go unassigned.

3. Exonic bins: An exonic bin is a concept introduced by DEXSeq (Anders, et al., 2012). It represents regions that are exonic in at least one annotated isoform of a given gene. Boundaries between exonic bins are either splice junctions are regions where the set of isoforms in which the region is exonic changes. A custom python and shell script is first called to flatten the provided annotation in the style of DEXSeq. Exonic bin assignment is done with featureCounts, providing this flattened annotation. The parameters used for this by default are: `-R CORE -f -g exon_id -t exonic_part -O -M`. If paired-end reads are provided, the following is automatically added to the featureCounts parameters: `-p –countReadPairs`. The output CORE file is used in all downstream steps of the pipeline. Unlike with Genes (exonic only), reads overlapping intronic regions will go assigned. Thus, it is advised to combine this assignment with Genes (exonic only) to filter out or disambiguate reads from likely pre-mRNA as necessary.
4. Exon-exon junctions: To perform this feature assignment, users must either provide bam files with STAR's unique jI and jM tags present, or they must provide fastq files and use STAR with fastq2EZbakR. A custom Python script processes the jI tag to denote the set of junction starts and end each read contains. STAR's custom jM tag controls filtering of novel, unannotated junctions discovered via STAR's 2-pass mapping mode (Veeneman, et al., 2016).
5. Transcript equivalence classes (TEC): TECs represent the set of annotated transcript isoforms with which a read is fully compatible. To perform this feature feature assignment, users must provide fastq files and use STAR to align their reads in fastq2EZbakR. A custom Python script is used to parse the resulting transcriptome aligned bam produced by STAR.

The `-M` flag (assign multi-mapping reads) is included in all featureCounts calls by default, as fastq2EZbakR filters out non-primary alignments prior to feature assignment. The `-M` flag ensures that the primary alignment of multi-mapping reads is assigned.

*Processing and analyzing real data*

Real NR-seq data (Figures 3C, S2, S3, and S6) was obtained from a recent publication performing an extensive suite of subcellular fractionation NR-seq experiments (Ietswaart, et al., 2024). The human (K562) total RNA, nuclear RNA, and cytoplasmic RNA samples were processed using fastq2EZbakR and all of the feature assignment strategies (users can select any subset of these in practice, and only gene-anywhere and gene-exonic were used in this manuscript). For Figures 3C and S2, reads were assigned to features in the full, unfiltered Ensembl annotation (GRCh38.108) (Harrison, et al., 2024). For Figures S3 and S6, reads were assigned to features in the same annotation, but filtered for only support level 1 and 2 isoforms, to improve the accuracy of exonic vs. intronic assignment.

For Figures 3C and S2, EZbakR's hierarchical mixture model was fit to the 2 hour $s^4U$ label time total RNA data with EstimateFractions() to the reads assigned to both a gene-anywhere and gene-exonic feature (i.e., intronic reads were filtered out). Default parameters were used.

For Figure S3, the following steps were followed to fit the pre-mRNA processing model to total RNA NR-seq data:

1. The fraction labeled for each gene (exonic and intronic reads separately; XF and GF columns of cB) was estimated with EstimateFractions, using -$s^4U$ control data to infer a single $p_{\text{unlabeled}}$ for all samples.
2. Dropout was corrected for using a method previously implemented in bakR (mathematical details here: https://simonlabcode.github.io/bakR/articles/Dropout.html), and similar to the method implemented in grandR (Rummel, et al., 2023).
3. Replicate logit fraction labeled estimates were averaged and length normalized read counts computed using AverageAndRegularize() with the formula ~tl (meaning that averaging was performed for each label

time (tl) independently). Intronic and exonic feature lengths were derived from the annotation file used for feature assignment, and custom R scripting that made use of the GenomicFeatures library (Lawrence, et al., 2013).

4. EZDynamics was used, with default settings, to fit the model of RNA maturation presented in Figure 5B. Reads mapping to definitively intronic regions were assumed to represent the population labeled P (pre-mRNA) in that model, and those mapping to definitively exonic regions were assumed to represent the population labeled M (mature mRNA) in that model. Technically, the latter is a combination of definitively mature and ambiguously mature or premature mRNA reads, but adding this distinction through a splice junction presence Boolean flag had limited impact on model fit.

For Figure S6, total, cytoplasmic, and nuclear NR-seq data was analyzed following the same steps were followed, except that:

1. Exclusively exonic read were analyzed.
2. The three datasets were combined into a single arrow dataset file system and analyzed using EZbakR's optional Apache arrow backend to reduce RAM usage (only relevant for step 1, fraction estimation) (Lentner, 2019).
3. The formula for AverageAndRegularize() was ~tl:compartment, where compartment describes whether or not the data came from total, cytoplasmic, or nuclear RNA. Thus, averaging was performed for each unique combination of label time and compartment.
4. The model fit by EZDynamics was that schematized in Figure 5C. Nuclear RNA data was assumed to represent the population labeled N (nuclear RNA), cytoplasmic RNA data was assumed to represent the population labeled C (cytoplasmic RNA), and total RNA data was assumed to represent the sum of N and C.
5. PUNDs were identified as those with low uncertainty (< 0.3) $\log(k_{Ndeg})$ estimates, all of which were similar in magnitude to the gene's $\log(k_{exp})$ estimate, representing significant amounts of putative nuclear degradation.

*Generalized mixture modeling in EZbakR*

As mentioned in Methods, mutational mixture modeling in EZbakR is performed in two steps:

1. Mutation rates in labeled and unlabeled reads ($p_{\text{labeled}}$ and $p_{\text{unlabeled}}$) for all types of mutations being modeled are estimated. This is done by fitting a two-component mixture model for each mutation type to all reads from a given sample. Fitting is done for each mutation type independently at this stage. If users have control samples lacking any labeling, they can optionally choose to use a single $p_{\text{unlabeled}}$ derived from these samples. This can improve the stability of $p_{\text{labeled}}$ estimates in some settings.
2. The generalized mixture model parameters are estimated via the method of prior-penalized maximum likelihood.

The generalized likelihood function for a single sequencing read can be formalized as such:

$$\mathrm{L}(\boldsymbol{\theta}, \boldsymbol{p}) = \sum_{i}^{\mathrm{P}} \theta_i * \prod_{j}^{\mathrm{M}} \mathrm{Bin}(\mathrm{nM}_j, \mathrm{nN}_j, p_{i,j})$$

$$\mathrm{Bin}(\mathrm{nM}, \mathrm{nN}, p) = \frac{\mathrm{nM}!}{\mathrm{nM}! * (\mathrm{nN} - \mathrm{nM})!} * p^{\mathrm{nM}} * (1-p)^{\mathrm{nN}-\mathrm{nM}}$$

P = Number of mutational populations present

M = Number of mutation types being modeled (e.g., T-to-C and G-to-A)

$\boldsymbol{\theta}$ = P x 1 vector of population fractions

$\boldsymbol{p}$ = P x M matrix of mutation rates. Element i, j represents rate of mutations of type j (i.e., high or low) in population i

nN = Number of mutable nucleotides of a particular type in reference sequence with which read overlaps

nM = Number of mutations of a particular type

Bin(nM, nN, p) = binomial distribution likelihood for nM mutations in nN mutable nuleotides and probability of mutation p.

For P = 1 and M = 1, this reduces to the standard two-component mixture model fit in step 1:

$$\mathrm{L}(\theta, p_{\mathrm{labeled}}, p_{\mathrm{unlabeled}}) = \theta_{\mathrm{labeled}} * \mathrm{Bin}(\mathrm{nM}, \mathrm{nN}, p_{\mathrm{labeled}}) + (1 - \theta_{\mathrm{labeled}}) * \mathrm{Bin}(\mathrm{nM}, \mathrm{nN}, p_{\mathrm{unlabeled}})$$

The log of these likelihoods are maximized with R's optim() function using the L-BFGS-B method. The actual likelihood maximized reparametrizes all elements of $\boldsymbol{\theta}$ and $\boldsymbol{p}$ in terms of their log-odds (i.e., logits). Weakly regularizing normal priors are used for all estimated parameters. These priors were chosen through prior predictive simulations aimed at capturing the standard range of $\boldsymbol{\theta}$ and $\boldsymbol{p}$ values seen across a wide array of distinct datasets. Users are able to adjust these priors as they see necessary. Uncertainties for all parameter estimates are obtained from the Hessian matrix provided as optional output by optim(). More specifically, a parameter's uncertainty is estimated to be the square root of the relevant diagonal element of the inverted Hessian.

*Generalized linear dynamical systems modeling in EZbakR*

As described in Methods, generalized linear dynamical systems modeling in EZbakR is performed with the EZDynamics() function. It can take as input either sample-specific fraction labeled estimates (output of EZbakR's EstimateFractions() function) or condition-wide averages obtained from the EZbakR generalized linear model fit (AverageAndRegularize() function). Both are compatible with modeling of pre-mRNA dynamics, but modeling of RNA flow between subcellular compartments is only compatible with the latter. This is because modeling of subcellular compartment flow requires integrating across multiple independent samples (i.e., data from different subcellular fractions), so there is no way to estimate all kinetic parameters of such models for individual samples. The input to EZDynamics() is a matrix representation of graphical models akin to those in Figure 5. Documentation regarding model specification can be found here: https://isaacvock.github.io/EZbakR/articles/EZDynamics.html. Other input includes information about how the measured features relate to the modeled features. For example, if fitting the subcellular fractionation model shown in Figure 5, you will likely have whole cell data. The RNA from this sample corresponds to the sum of N (nuclear RNA) and C (cytoplasmic RNA), and this fact must be conveyed to EZDynamics().

EZDynamics() then uses the method of maximum likelihood to estimate kinetic parameters of your model. This is done by modeling both the fraction labeled and the read counts (if normalization is possible) for a given feature. If sample-specific fraction labeled estimates are provided as input, then the following model is assumed:

$$\mathrm{logit}(\theta) \sim \mathrm{Normal}(\mathrm{f}(\mathbf{k}), \theta_{\mathrm{uncert}})$$

$$\text{Read counts} \sim \mathrm{Poisson}(\mathrm{g}(\mathbf{k})*\mathrm{scale})$$

$\theta$ = sample and feature-specific fraction labeled estimate

$\mathbf{k}$ = vector of kinetic parameter estimates

f() and g() = functions mapping kinetic parameters to expected fraction labeled and abundance, respectively

$\theta_{\mathrm{uncert}}$ = Uncertainty in logit($\theta$)

scale = Normalization scale factor; default is TMM strategy similar to that used in DESeq2

If condition-wide averages are provided, the following model is assumed:

$$\mathrm{logit}(\theta_{\mathrm{avg}}) \sim \mathrm{Normal}(\mathrm{f}(\mathbf{k}), \sigma_{\mathrm{post}})$$

$$\log_{10}(\mathrm{Avg.\ read\ coverage}) \sim \mathrm{Normal}(\mathrm{g}(\mathbf{k}) * \mathrm{scale}, \mathrm{C}_{\mathrm{uncert}})$$

$\theta_{\mathrm{avg}}$ = condition-wide fraction labeled average

$\sigma_{\mathrm{post}} = \mathrm{logit}(\theta_{\mathrm{avg}})$ posterior standard deviation

$\mathrm{C}_{\mathrm{uncert}}$ = delta approximation of the expected $\log_{10}$(coverage) standard deviation assuming reads ~ Poisson

f(**k**) and g(**k**) are inferred from the analytic solution to the linear system of ODEs implied by a given graph. The analytic solution is derived from a matrix representation of the system of ODEs, as such (Ricardo, 2020):

$$\frac{\mathrm{d}\mathbf{r}}{\mathrm{dt}} = \mathbf{M} * \mathbf{r}(\mathrm{t})$$

**M** = N x N matrix, where N = number of RNA species being modeled

**r**(t) = N x 1 vector of functions of modeled RNA species' levels over time (t)

$$\mathbf{r}(\mathrm{t}) = \sum_{i}^{\mathrm{N}} v_i * e^{\lambda_i * \mathrm{t}}$$

$v_i$ = ith eigenvector of **M**

$\lambda_i$ = ith eigenvalue of **M**

Technically, **r**(t) as presented above is only the general solution in the case where **M** has N unique eigenvalues. In this context, this is roughly equivalent to all kinetic parameter estimates being distinct. As these estimates should be thought of as continuous random variables though, the probability of any two eigenvalues being equal is 0. Thus, we can assume the degenerate eigenvalue edge case is never relevant. EZDynamics() avoids this edge case computationally by introducing a small amount of noise to parameter estimates if they are equal in any given iteration of the optimization procedure.

Theoretical identifiability is assessed via first checking that the number of estimated parameters is less than or equal to twice the number of measured species (since each measured RNA species yields a fraction labeled and read count estimate). Practical identifiability can be assessed via the Hessian, with practically non-identifiable parameter estimates yielding very high uncertainties for that parameter (square root of the relevant diagonal element of the inverse of the Hessian) (Rodriguez-Fernandez, et al., 2006a; Rodriguez-Fernandez, et al., 2006b). Parameters associated with non-productive routes along a forked path represent a particularly common case of practical unidentifiability (e.g., the model in Figure 5C, where pre-mRNA can either be processed into mRNA or degraded). Figure 5D shows that the uncertainty estimates derived from the Hessian matrix provides a good metric for when such parameters are practically non-identifiable. The intuition for why estimation of these types of parameters in particular are challenging to estimate can be developed via considering the 3 possible cases (see Figure 5D, right for evidence of the intuitive explanations described below):

1. If the rate constant for the unproductive route is much faster than that of any of the productive routes, it means that the levels of downstream RNA species will be very low. This will typically yield low read coverage and thus low confidence estimates of the fraction labeled for these species. Thus, in this case, it is difficult to determine if the low abundance of these species is due to the large value of the unproductive

route's rate constant, or high amounts of turnover of this RNA species. This leads to practical unidentifiability of the unproductive route's rate constant.

2. If the rate constant for the unproductive route is much slower than that of any of the productive routes, it means that its exact value has little impact on the measured quantities (e.g., read counts and fraction labeled's). For example, the abundance of premature RNA in the model in Figure S4A is a function of the sum of the productive and unproductive rate constants (Pss = $k_{syn}/(k_p + k_{pdeg})$; $\theta_P = 1 - e^{-(k_p + k_{pdeg})*tl}$). When the two rate constants differ by an order of magnitude or more, this sum is dominated by the larger term and is not strongly impacted by changes in the smaller term. Thus, if the unproductive route rate constant is much smaller, it is practically unidentifiable.
3. If the two rate constants on a forked path are of similar orders of magnitude, then both are typically practically identifiable.

Estimating scale factors for multi-compartment modeling can be done in one of three ways:

1. Spike-in normalization. Scale factors should be calculated by users and provided to EZbakR.
2. Fraction labeled modeling. The fraction labeled estimates (which are internally normalized) can be used to estimate all but the RNA synthesis parameter, if such an estimation strategy is identifiable. Scale factors can then be inferred from the set of non-synthesis parameter estimates, by identifying the factors by which absolute RNA abundances are expected to differ across the compartments (since the synthesis rate is the same for all RNA species from a given gene). A downside of this strategy is that it limits the complexity of identifiable models, and typically yields lower confidence parameter estimates (Figure S4, top row).
3. Fraction labeled mixing model. If users have data for individual subcellular compartments as well as data for a sufficient number of combinations of these compartments, then scale factors can be estimated from the differences in overall fraction of reads that are labeled in each compartment or combination of compartments, for a given metabolic labeling time. This strategy doesn't constrain the space of identifiable models that can be fit in the ways that option 2 does. For example, if users have nuclear fraction, cytoplasmic fraction, and whole cell data, then normalization factors can be estimated by solving the following linear system of equations and using the inferred ratio of absolute molecular abundances to derive scale factors for each compartment:

$$\theta_{\mathrm{wc}} = \theta_{\mathrm{nuc}} * \frac{\mathrm{Nss}}{\mathrm{Css} + \mathrm{Nss}} + \theta_{\mathrm{cyto}} * \left(1 - \frac{\mathrm{Nss}}{\mathrm{Css} + \mathrm{Nss}}\right)$$

$$\mathrm{s} = \frac{\mathrm{Nss}}{\mathrm{Css} + \mathrm{Nss}}$$

$$\text{scale factors} = 1 \text{ for Nuclear data (arbitrary)}; \frac{1 - \mathrm{s}}{\mathrm{s}} \text{ for cytoplasmic data}; 1 + \frac{1 - \mathrm{s}}{\mathrm{s}} \text{ for whole cell data}$$

Nss = steady-state molecular abundance of nuclear RNA

Css = steady-state molecular abundance of cytoplasmic RNA

$\theta_{\mathrm{nuc}}$ = total fraction of reads per kilobase (RPK) labeled in nuclear data

$\theta_{\mathrm{cyto}}$ = total fraction of reads per kilobase (RPK) labeled in cytoplasmic data

$\theta_{\mathrm{wc}}$ = total fraction of reads per kilobase (RPK) labeled in whole cell data

We note that option 3 follows from the same model used to derive a normalization scheme proposed in a recent preprint, where the idea of modeling the whole cell data as the weighted sum of nuclear and cytoplasmic data was first introduced (Steinbrecht, et al., 2024). That work assumed that scale factors could be derived from the relative estimated transcript per millions (TPMs) of cytoplasmic, nuclear and whole cell RNA. While TPMs normalize for library size and transcript length, they do not contain information about absolute abundance in a way that is necessary for robust normalization in this setting. Thus, we use the length-adjusted total fraction labeled, an internally normalized quantity that allows for accurate scale factor inference in this case. See Supplemental Methods for further details regarding normalization, and a discussion of theoretical and practical identifiability of certain ODE systems.

To derive the exact form of the scale factors in scenario 3, consider a simple model for the expected read counts of a given feature in a given RNA-seq sample:

$$\mathrm{E[read\ counts\ for\ feature\ i\ in\ nuclear\ fraction]} = \frac{m_{i,N} * L_i}{\sum_{j=1}^{N} m_{j,N} * L_j} * L_i * R_N = \frac{m_{i,N} * L_i}{\mathrm{Nss}} * R_N$$

$$\mathrm{E[read\ counts\ for\ feature\ i\ in\ nuclear\ fraction]} = \frac{m_{i,C} * L_i}{\sum_{j=1}^{N} m_{j,C} * L_j} * L_i * R_C = \frac{m_{i,C} * L_i}{\mathrm{Css}} * R_C$$

$$\mathrm{E[read\ counts\ for\ feature\ i\ in\ whole\ cell\ data]} = \frac{m_{i,WC} * L_i}{\sum_{j=1}^{N} m_{j,WC} * L_j} * R_{WC} = \frac{m_{i,WC} * L_i}{\mathrm{WCss}} * R_{WC}$$

m = number of molecules from a given feature

N = total number of features sequenced

$L_i$ = effective length of feature i

R = total number of reads in a given sample

The expected read count is not only influenced by the absolute molecular levels of a particular feature (numerator of all expressions), but also by the total molecule content of a given sample (denominator of all expressions), the length of the feature (L), and the sequencing depth of a given sample (R). The latter two can be easily dealt with via standard RPKM or TPM normalization. In order to put read counts on a scale such that differences in read counts are a function of differences in absolute molecular levels, we need to identify scale factors that if we multiply read counts in each sample by, eliminate dependence on the unique denominators. For example, if we knew the ratio of Nss to Css and WCss (Css/Nss and WCss/Nss), we could multiply cytoplasmic and whole cell read counts by these ratios to yield normalized read counts with the same dependence on absolute molecular levels:

$$\frac{\mathrm{E[read\ counts\ for\ feature\ i\ in\ nuclear\ fraction]}}{R_N * L_i} * 1 = \frac{m_{i,N}}{\mathrm{Nss}}$$

$$\frac{\mathrm{E[read\ counts\ for\ feature\ i\ in\ nuclear\ fraction]}}{R_C * L_i} * \frac{\mathrm{Css}}{\mathrm{Nss}} = \frac{m_{i,C}}{\mathrm{Nss}}$$

$$\frac{\mathrm{E[read\ counts\ for\ feature\ i\ in\ whole\ cell\ data]}}{R_{WC} * L_i} * \frac{\mathrm{WCss}}{\mathrm{Nss}} = \frac{m_{i,WC}}{\mathrm{Nss}}$$

The length normalized global fraction labeled (length normalized fraction of all reads from RNA synthesized during the label time; estimated as $\mathrm{sum}(\theta_i * \mathrm{RPKM_i})/\mathrm{sum}(\mathrm{RPKM_i})$) in each subcellular fraction is a function of these ratios, since:

$$\theta_{\mathrm{wc}} = \frac{\theta_{\mathrm{nuc}} * \mathrm{Nss} + \theta_{\mathrm{cyto}} * \mathrm{Css}}{\mathrm{Nss} + \mathrm{Css}}$$

$$\theta_{\mathrm{wc}} = \theta_{\mathrm{nuc}} * \frac{\mathrm{Nss}}{\mathrm{Css} + \mathrm{Nss}} + \theta_{\mathrm{cyto}} * \left(1 - \frac{\mathrm{Nss}}{\mathrm{Css} + \mathrm{Nss}}\right)$$

$$\theta_{\mathrm{wc}} = \theta_{\mathrm{nuc}} * \mathrm{s} + \theta_{\mathrm{cyto}} * (1 - \mathrm{s})$$

From this linear equation, we can derive an expression for s:

$$\mathrm{s} = \frac{\theta_{\mathrm{wc}} - \theta_{\mathrm{cyto}}}{\theta_{\mathrm{nuc}} - \theta_{\mathrm{cyto}}}$$

The ratio of Nss to Css can then be derived from s as follows:

$$\mathrm{s} = \frac{\mathrm{Nss}}{\mathrm{Css} + \mathrm{Nss}}$$

$$\mathrm{s} * (\mathrm{Css} + \mathrm{Nss}) = \mathrm{Nss}$$

$$\mathrm{s} * \mathrm{Css} = \mathrm{Nss} * (1 - \mathrm{s})$$

$$\frac{\mathrm{Css}}{\mathrm{Nss}} = \frac{(1 - \mathrm{s})}{\mathrm{s}}$$

This is exactly the scale factor for cytoplasmic fraction data presented above. To derive the scale factor for whole cell data, note that WCss = Nss + Css:

$$\mathrm{WCss} = \mathrm{Nss} + \mathrm{Css}$$

$$\mathrm{WCss} = \mathrm{Nss} + \mathrm{Nss} * \frac{(1 - \mathrm{s})}{\mathrm{s}}$$

$$\mathrm{WCss} = \mathrm{Nss} * \left(1 + \frac{(1 - \mathrm{s})}{\mathrm{s}}\right)$$

$$\frac{\mathrm{WCss}}{\mathrm{Nss}} = 1 + \frac{(1 - \mathrm{s})}{s}$$

In EZbakR, scale factors are estimated based on average global fraction labeled for a set of compartments for a given label time. This is done through maximum likelihood estimation of a generalization of the procedure described above. We then use the lowest uncertainty set of scale factors (estimated using the Hessian) across the label times to get the final compartment-specific scale factor. If label times used for data from one subcellular fraction differ from all of those used in the other fractions, then this normalization strategy cannot be used. The generalization of this procedure is formalized below:

$$\vec{\theta} \sim \mathrm{Normal}((\mathrm{M} * (\overrightarrow{\mathrm{v}_1} \otimes \overrightarrow{\mathrm{v}_2})) \oslash (\mathrm{M} * \overrightarrow{\mathrm{v}_2}), \sigma)$$

$\vec{\theta}$ = N x 1 vector of global fraction labeled estimates in each sample

M = N x S matrix specifying the modeled RNA species present in each sample

N = number of samples

S = number of modeled RNA species

$\overrightarrow{\mathrm{v}_1}$ = S x 1 vector of RNA species global length normalized fraction labeled estimates

$\overrightarrow{\mathrm{v}_2}$ = S x 1 vector of relative molecular abundances of modeled RNA species; one element is set to 1 by dividing all other elements by its value.

$\otimes$ = element-wise multiplication (Hadamard multiplication)

$\oslash$ = element-wise division (Hadamard division)

σ = uncertainty in global fraction labeled estimate

In this case, a *sample* typically represents the average estimates for multiple replicates of a given label time from a given population of RNA (e.g., nuclear RNA). The unknown parameters (non-trivial elements of $\overrightarrow{v_1}$ and $\overrightarrow{v_2}$; the trivial element is the 1 in $\overrightarrow{v_2}$) are estimated via the method of maximum likelihood, and uncertainties are estimated using the Hessian. In the total-cytoplasmic-nuclear RNA case derived above, the terms specified in the generalized model could look like:

$$\vec{\theta} = [\theta_{\text{nuc}}, \theta_{\text{cyto}}, \theta_{\text{wc}}]$$

$$\text{M} = \begin{bmatrix} 1 & 0 \\ 0 & 1 \\ 1 & 1 \end{bmatrix}$$

$$\overrightarrow{v_1} = [\theta_{\text{N}}, \theta_{\text{C}}]$$

$$\overrightarrow{v_2} = [1, \text{Css}/\text{Nss}]$$

$\theta_{\text{N}}$ = true proportion of nuclear RNA that is labeled; estimated

$\theta_{\text{C}}$ = true proportion of cytoplasmic RNA that is labeled

Css = Molecular abundance of cytoplasmic RNA

Nss = Molecular abundance of nuclear RNA

The first column in M represents whether nuclear RNA is present in a given sample, and the second column represents whether cytoplasmic RNA is present. In this case, there is a single label time of data from nuclear (first row), cytoplasmic (second row), and total RNA (third row). M is inferred automatically from standard input to EZDynamics(). Scale factors for each sample are estimated as:

$$\text{scale factors} = \text{M} * \overrightarrow{v_2}$$

In the total-cytoplasmic-nuclear case, this yields the same expression for the scale factors derived above.

*Generalized linear modeling in EZbakR*

EZbakR's AverageAndRegularize() function is used to fit a generalized linear model of a user's fraction labeled or kinetic parameter estimate data. Parameter standard error estimates are then regularized using a hierarchical modeling strategy similar to that introduced in bakR (Vock and Simon 2022). The EZbakR input data object (an EZbakRData object) includes a so-called metadf table, which can contain any number of factors describing elements of each sample (see example in Figure 6). These factors can be specified in a formula object passed to AverageAndRegularize(), which will specify the linear model to fit to one's data. A unique aspect of EZbakR's generalized linear model is that it can allow for heteroskedasticity through a number of different ways. 1) If a user's model is such that each parameter is estimated from a non-overlapping set of samples, then standard deviations are estimated for each of these sample sets independently or, 2) users can specify a set of factors by which to group samples, and standard deviations are estimated in these sample sets and these standard deviations are used to estimate parameter standard errors. More specifically, group standard deviations are plugged into a diagonal matrix $\mathbf{\Omega}$ where the ith diagonal element is the ith data point's group standard deviation and parameter standard errors are calculated as usual for linear regression:

$$\text{standard error} = (\mathbf{X}^{\text{T}} * \mathbf{X})^{-1} * \mathbf{X}^{\text{T}} * \mathbf{\Omega} * \mathbf{X} * (\mathbf{X}^{\text{T}} * \mathbf{X})^{-1}$$

**X** = design matrix

EZbakR improves upon bakR by balancing extra conservativeness in several steps with a more highly powered statistical testing scheme. In particular, the following changes to the regularization scheme were made:

1. Sample-specific parameter uncertainties were used to generate conservative estimates of feature-specific replicate variabilities. In addition, a small floor is set to ensure that standard deviation estimates are never below a certain level, for the same reason.
2. Condition-wide replicate variabilities were regressed against both read coverage, and either:
    a. |logit(θ)| when modeling fraction labeled. This captures the fact that estimates are best around a logit(θ) of 0 and get worse for more extreme fraction labeled.
    b. log(kdeg) when modeling log degradation rate constants. At first, we considered a strategy similar to the fraction labeled modeling, but found that the non-linear transformation of fraction lbaeled to log(kdeg) yields complicated relationships between the two parameter's uncertainties. Empirically, we found that agreement between a fully rigorous MCMC sampling approach and EZbakR was significantly improved by just regressing the value of the log kinetic parameter.
    c. Only coverage in other cases.
3. Features with replicate variabilities below the inferred trend had their replicate variabilities set equal to that predicted by the trend. This helps limit underestimation of parameter variance. Features with above-trend replicate variabilities had their replicate variabilities regularized with a Normal prior Normal likelihood Bayesian model as in bakR.
4. P-values are calculated using a Wald test (less conservative) rather than a moderated t-test. The conservativeness introduced by the above changes allowed for this change without sacrificing FDR control (Figure 6 C-E).

*Generating simulated data*

Simulated data used in this study were generated with simulation functions implemented in EZbakR. Multi-label data was simulated with EZbakR's SimulateMultiLabel() function. Data with feature-to-feature $p_{\text{labeled}}$ variation was simulated with EZbakR's SimulateOneRep() function. Data for all dynamical systems models tested in Figure 4 were simulated with EZbakR's SimulateDynamics() function.

SimulateOneRep() details:

- Data for 5000 features was simulated for analyses in Figure 4
- Reads of 200 nucleotides in total length (e.g., PE100 reads) were simulated.
- The number of Us in each read was drawn from a binomial distribution with number of trials equal to the read length and probability of each nucleotide being a U equal to 0.25
- A feature's degradation rate constant was drawn from a log-normal distribution with log-mean = -1.9 and standard deviation on the log scale of 0.7. This yields a kdeg distribution reminiscent of that seen in human cell line data.
- The expected fraction of reads that are labeled was then 1 – exp(-kdeg*2); i.e., a 2 hour label time was simulated. The actual labeled vs. unlabeled identify of a read was simulated using a Bernoulli distribution with probability the read is labeled being the relevant feature's fraction labeled.
- Figure 4 investigates EZbakR's ability to estimate feature-specific $p_{\text{labeled}}$. Thus, each simulated feature was also simulated a unique $p_{\text{labeled}}$ value. These were drawn from a logit-normal distribution with logit-mean of -2.5 and standard deviation on the logit scale of 0.1 (Figure 4B left).
- The number of mutations in a sequencing read was drawn from a binomial distribution with number of trials equal to number of Us, and the probability of a mutation being either the background mutation rate

of 0.002 (if read was unlabeled) or the feature-specific $p_{labeled}$ if the read was from labeled RNA. We clarify that a read from labeled RNA can thus have 0 mutations.

SimulateMultiLabel() details:

- TILAC data was simulated, which means that a simplex vector of length three needs to be simulated for each feature (the fraction of reads that are $s^4U$ labeled, $s^6G$ labeled, and unlabeled; simplex vector as these values must sum to 1).
- The simplex vector was drawn from a Dirichlet distribution, with the vector of alphas drawn from a Uniform(3, 6) distribution.
- Counts for Us and Gs in a read were drawn from a multinomial distribution, with each nucleotide in a read having an equal probability of one of the four nucleotides.
- Mutational data was simulated as in SimulateOneRep(), with all features having the same labeled and unlabeled T-to-C and G-to-A mutation rates.

SimulateDynamics details:

- The set of feature-specific kinetic parameters simulated for a given model was drawn from a log-normal distribution.
    - For the 0 → P → M → 0 model, the log-mean of the pre-mRNA processing parameter was -0.3, and the log-mean of the RNA degradation parameter was -2.
    - For the nuclear + cytoplasmic RNA model (Figure 5C), the log-mean of the nuclear export parameter was -0.3, the log-mean of nuclear degradation parameter was -1 (so typically slower than export, meaning that most RNAs don't undergo significant nuclear degradation), and the log-mean of the cytoplasmic RNA degradation parameter was -2. These parameters were the same for the simulation without nuclear degradation (except obviously the lack of a nuclear degradation parameter) (Figure S4).
    - For the more complex model with pre- and mature RNA nuclear and cytoplasmic dynamics, the log-means were arbitrarily set to the following range of different values to challenge the model with different ranges of kinetic parameters:
        - -0.3 for nuclear pre-mRNA processing
        - -0.725 for nuclear pre-mRNA export
        - -1.150 for nuclear mature RNA export
        - -1.575 for cytoplasmic pre-mRNA processing
        - -2 for cytoplasmic mature RNA degradation
    - The log-mean of the RNA synthesis rate was set to 1 in all simulations (choice is arbitrary)
    - A log-scale standard deviation of 0.4 was used for all simulations
- The analytic solution for a given feature's new and old RNA levels was used to determine expected fraction labeled. These were passed to SimulateOneRep()
    - Two replicates of one hour label times, and two replicates of three hour label times were simulated in all cases.
